# The inflammasome sensor NLRP3 interacts with REV7 to maintain genome integrity through homologous recombination

**DOI:** 10.1101/2024.11.28.625806

**Authors:** Delphine Burlet, Md Muntaz Khan, Sabine Hacot, Hannes Buthmann, Léa Bardoulet, Anne-Laure Huber, Julie Gorry, Bernard S Lopez, Yohann Couté, Matthias Geyer, Agnès Tissier, Virginie Petrilli

## Abstract

DNA double strand break (DSB) is a highly toxic lesion that can generate genome instability, a major source of tumorigenesis. DSBs are mainly repaired by non-homologous end joining (NHEJ) or homologous recombination (HR). The selection of the DSB repair pathway primarily depends on the DNA resection of the DSB ends. Indeed, HR is initiated by resection at the DSB generating 3’ single stranded extension. The shieldin complex prevents resection fostering DSB repair toward NHEJ. Here, we reveal that the inflammasome sensor NLRP3 facilitates DNA end resection to promote the HR pathway in an inflammasome-independent manner. Strikingly, NLRP3 silencing decreases HR efficiency, as evidenced by RAD51 foci and functional HR assays. Mechanistically, we describe that NLRP3 interacts with REV7, a subunit of the shieldin complex, and its depletion increases REV7 recruitment to IR-induced DSBs. Similar to cancer cells harboring HR mutated genes, we find that NLRP3 deficient cells are sensitive to PARP inhibitors (PARPi) and exhibit an epistatic relationship with BRCA1 deficiency. Remarkably, loss of REV7 in NLRP3-depleted cells induces PARPi resistance by restoring HR. This study unravels the crucial role of the innate immune receptor NLRP3 in regulating the selection of DSB repair pathways to maintain genome integrity.

## INTRODUCTION

Maintaining genomic integrity is critical for tissue homeostasis. Genomic DNA is constantly exposed to stresses of exogenous and endogenous origin. DNA damage is detected by a sensing machinery called the DNA damage response (DDR), which promotes DNA repair^1^. DNA double-strand breaks (DSBs) are the most toxic lesions as defects in their repair result in increased chromosomal instability, premature aging and tumorigenesis^2^. These breaks are repaired by two major pathways: non homologous end joining (NHEJ) and homologous recombination (HR). The HR pathway is considered to be the most faithful DDR, since it uses homologous sequences from the sister chromatid as a repair template. Cancer cells harboring mutations in genes affecting this pathway, such as *BRCA1/2*, are HR deficient (HRD) and vulnerable to PARP inhibitors (PARPi)^3^. Although PARPi have improved patient survival, resistance mechanisms have emerged^4,5^. One of them, in the *BRCA1* mutated background, relies on the inactivation of the shieldin complex^6–8^, which is composed of the SHLD-1,-2,-3 and REV7 proteins^9,10^. This complex acts downstream of 53BP1 and favors NHEJ by inhibiting DNA resection, a key step in HR^6,10–13^. Assembly and disassembly of this complex is regulated by TRIP13 that entraps REV7 to decrease its dimerizing ability^14,15^.

NLRP3 is a pattern recognition receptor (PRR) that fosters the formation of the inflammasome complex in response to pathogen- or damage-associated molecular patterns (PAMPs or DAMPs), inducing the production of mature IL-1β and IL-18, and pyroptosis^16,17^. NLRP3 plays a critical role in the maintenance of tissue homeostasis, by sensing a wide range of stresses^18^. Though this PRR is mostly considered as a cytosolic protein, nuclear and inflammasome-independent functions are emerging. In the context of genome stability, NLRP3 was recently reported to interact with ATM and to enhance its activation in response to DNA DSBs^19,20^. Whether NLRP3 controls the DSB repair machinery remains unknown.

Here, we report that NLRP3 is located in close proximity to γH2AX foci, and demonstrate that it promotes the HR pathway by inhibiting REV7 to allow DNA end resection. HR deficiency is further supported by showing that NLRP3 expression is inversely correlated with the homologous recombination repair deficiency (HRD) score in breast cancers and by PARPi test sensitivity. Lastly, PARPi sensitivity is abolished by co-depletion of NLRP3/REV7, as previously demonstrated for BRCA1, re-enforcing the notion that NLRP3 and BRCA1 participate in the same functional pathway.

## RESULTS

### NLRP3 facilitates homologous recombination

To test whether NLRP3 is involved in DSB repair, we first induced DNA damage by ionizing radiation (IR) in control or NLRP3-depleted MDA-MB-231 cells (HR-proficient triple-negative breast cancer cell line expressing NLRP3^19^) and performed a comet assay. At the single-cell level, NLRP3 depletion by shRNA resulted in a significantly increase in tail moment suggesting an altered DNA repair capacity compared with control conditions after IR (Figure 1A). To strengthen this observation, DNA damage was induced using etoposide (Eto) treatment (inhibitor of topoisomerase II), and produced similar results in NLRP3-deficient cells (Sup Figure 1A), suggesting that NLRP3 plays a role in maintaining genome stability. Although, we previously showed that a fraction of NLRP3 is present in the nucleus^19^, whether it is recruited to the site of damage was not known. We next asked whether NLRP3 was located in the vicinity of DSBs by performing proximity-ligation assays (PLA) using antibodies directed against NLRP3 and γH2AX, a DSB marker. We specifically detected nuclear PLA signals only in response to IR treatment in MDA-MB-231 cells (Figure 1B and Sup Figure 1B) that were absent in shNLRP3- or siH2AX-treated cells, indicating that NLRP3 is in close proximity to DSBs following DNA damage.

**Figure 1.**
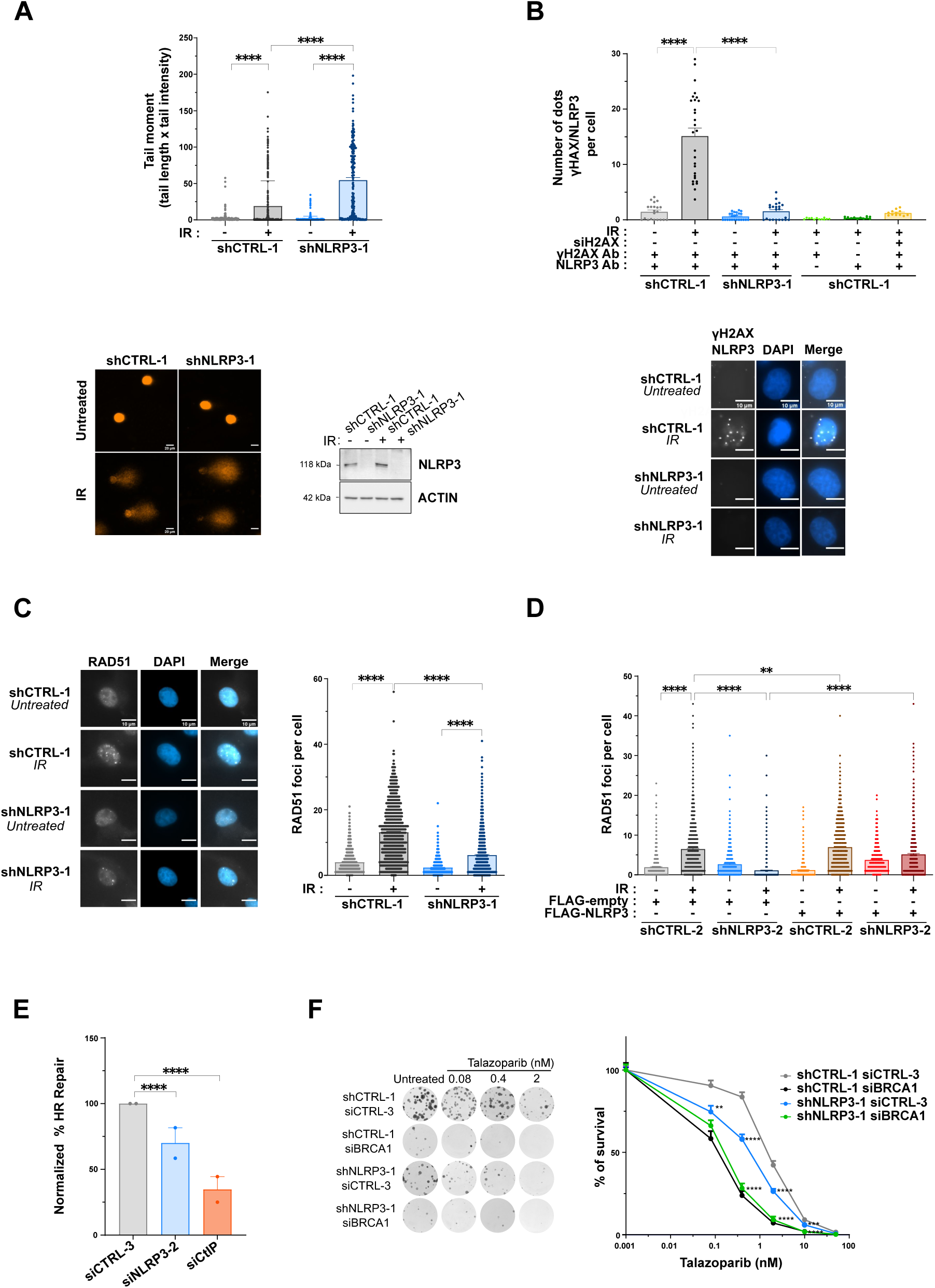
– NLRP3 fosters DNA damage repair through homologous recombination. **(A)** DNA damage was measured by comet assay in shCTRL-1 or shNLRP3-1 MDA-MB-231 cells and irradiated for 4 h at 5 Gy (IR). Representative images of comet tails (bottom panel), dot plot with comet tail moment (top panel), and control immunoblot demonstrating NLRP3 knockdown using ACTIN as a loading control (bottom panel). Scale bar: 20 µm. **(B)** Representative microscopy images of Proximity Ligation Assay (PLA) in white showing the interaction between γH2AX and NLRP3 (bottom panel). DAPI in blue. Scale bar: 10 µm. Quantification of the number of dots per cell in shCTRL-1 or shNLRP3-1 MDA-MB-231 cells 30 min after irradiation at 5 Gy (IR) (top panel). Additional controls were used to verify the specificity of the antibodies: cells were treated with siH2AX or labeling was performed using a single antibody against one of the targets (representative microscopy images Sup Figure 1B). **(C)** RAD51 foci in shCTRL-1 or shNLRP3-1 MDA-MB-231 cells treated with 5 Gy (IR), and fixed 6 h after treatment. Representative images of RAD51 foci (left panel) and dot plot with total number of foci per nucleus are shown (right panel). DAPI in blue. **(D)** shCTRL-2 or shNLRP3-2 MDA-MB-231 cells were transfected with a FLAG-empty or FLAG-NLRP3-expressing vector then treated with IR. Quantification of RAD51 foci per nucleus 6 h post IR (5 Gy) treatment foci is shown. **(E)** U-2 OS DR-GFP cells were co-transfected with siCTRL#3, siNLRP3#2 or siCtIP and HA-I-SceI-expressing plasmid, and the number of GFP+ cells was assessed by flow cytometry 72 h post-transfection. **(F)** Sensitivity of shCTRL-1 or shNLRP3-1 MDA-MB-231 cells transfected with control (CTRL-3) or BRCA1 siRNA to increasing doses of talazoparib was assessed using colony formation assay. Representative images of clones are shown (left panel). Percentage of surviving colonies was analyzed after 14 days (right panel). (A, B, C, F) shRNA-mediated NLRP3 knockdown was performed in MDA-MB-231. For (A, B, C, D), at least 100 cells were quantified. Mean and SEM for at least two (A, B, D, E) or three (C, F) independent replicates is shown. (A, B, C, D, F) The p-values correspond to a Mann Whitney’s test. (E) Chi2 squared test was performed. ns: nonsignificant, ** p < 0.01, *** p < 0.001, **** p < 0.0001.

To determine the mechanism underlying the defect in DSB repair in the absence of NLRP3, we next examined the formation of RAD51 and 53BP1 foci as markers of HR and NHEJ respectively in response to IR. We observed a strong decrease in RAD51 foci in cells lacking NLRP3 compared with controls and a small increase in 53BP1 foci suggesting a defect in HR (Figure 1C and Sup Figure 1C). This defect in RAD51 foci was also observed in response to Eto treatment (Sup Figure 1D). Consistently, the use of another shRNA targeting the 3’UTR sequence of NLRP3 together with the expression of exogenous NLRP3 rescued RAD51 foci formation demonstrating that NLRP3 acts on HR (Figure 1D and Sup Figure 1E).

To assess the role of NLRP3 on HR, we performed cell line-based reporter assays with U-2 OS cells containing one stable copy of the DR-GFP reporter cassette^21^ in which the expression of the enzyme HA-I-SceI induces a site-specific DSB. There were significantly fewer GFP+ cells following transfection with siNLRP3 and I-Sce1 compared to the siCTRL (Figure 1E et Sup Figure 1F). As a control, knock-down of CtIP, a nuclease of the resection step, results in decrease GFP+ cells^21^. Conversely, in cells displaying exogenous NLRP3 expression, an increase in the number of GFP+ cells was observed compared to the control, after I-SceI transfection (Sup Figure 1G). Taken together, these results show that NLRP3 positively regulates HR.

We next wondered whether NLRP3-depleted cells were sensitive to PARPi treatment, as *BRCA1*-deficient tumors are HR-deficient (HRD) and PARPi sensitive^3,22,23^. To test this hypothesis, we assessed the sensitivity of NLRP3-depleted MDA-MB-231 cells to talazoparib and olaparib, two PARPi with varying PARP trapping efficacies^24^. siBRCA1 was used as positive control, and as expected, BRCA1 depletion strongly sensitized cells to PARPi treatment (Figure 1F and Sup Figure 1H). Although milder than BRCA1 depletion, NLRP3 depletion resulted in significant sensitivity to both PARPi (Figure 1F and Sup Figure 1H). NLRP3 and BRCA1 co-depletion resulted in a similar level of sensitivity than BRCA1 alone, suggesting an epistatic relationship. Finally, U-2 OS cells expressing shNLRP3 were also sensitive to talazoparib (Sup Figure 1I), reinforcing the direct role of NLRP3 in this effect. Mutations in the *BRCA1* or *BRCA2* gene were originally associated with HRD. However, the notion of BRCAness phenotype has since been broadened and the HR-deficiency score is now calculated using the HRD and genome scarring score^25^. Here, we investigated the correlation between NLRP3 expression and the HRD score in breast cancers. Tumors expressing a high HRD score displayed lower levels of NLRP3, substantiating our findings that reduced NLRP3 expression is correlated with HR deficiency (Sup Figure 1J).

### NLRP3 is required for DNA end resection in HR

HR is initiated at DSBs by 5’ DNA end resection generating 3’ single stranded DNA (ssDNA) overhangs and bound by replication factor A (RPA) complex which can be visualized by immunofluorescence as RPA2 foci. Here, IR-induced RPA2 foci formation was strongly reduced upon NLRP3 depletion by respectively shRNA or siRNA in MDA-MB-231 (Figure 2A) and U-2 OS (Sup Figure 2A) compared to control. In line with these observations, the absence of NLRP3 also resulted in reduced etoposide-induced phosphorylation of RPA2 on Ser4/8, suggesting a defect in resection (Sup Figure 2B). To test whether NLRP3’s role on resection was an inflammasome-independent function, we quantified RPA2 foci formation in the presence of MCC950, a specific NLRP3 inflammasome inhibitor^26^. We found that MCC950 did not affect RPA2 foci recruitment (Sup Figure 2C). Given that MDA-MB-231 cells do not express the inflammasome core protein ASC^27^, which is required for inflammasome function (Sup Figure 2D), these results collectively support the notion that NLRP3, and not the inflammasome, is required for DSB repair via HR.

**Figure 2.**
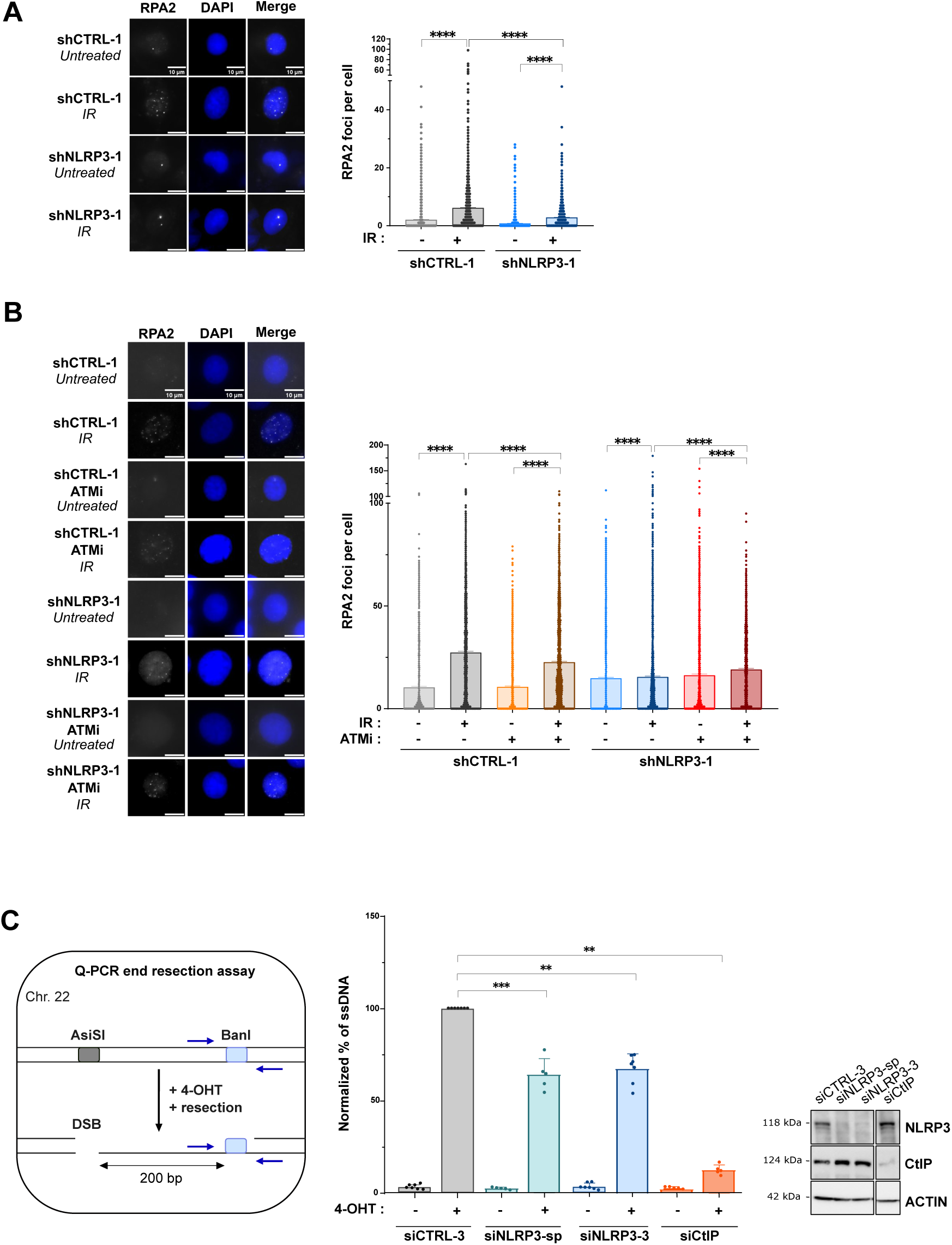
– NLRP3 depletion impairs DNA end resection. **(A)** RPA2-foci in shCTRL-1 or shNLRP3-1 MDA-MB-231 cells treated with 5 Gy (IR), fixed 6 h after treatment. Representative images of RPA2 foci (left panel) and dot plot with total number of foci are shown (right panel). DAPI in blue. Scale bar: 10 µm. **(B)** Quantification of RPA2 foci in shCTRL-1 or shNLRP3-1 MDA-MB-231 cells after IR in the presence of DMSO (-) or ATM inhibitor (KU55933) (+) (5 Gy, 6 h). Representative images of RPA2 foci and dot plot with total number of foci per cell are shown. DAPI in blue. Scale bar: 10 µm. **(C)** Schematic representation of PCR-based DNA end resection assay in DIvA cells (left panel). In vivo DSB was introduced at AsiSI site using 4-hydroxytamoxifen (4-OHT). DNA end resection was then monitored 200 bp away at BanI site on chromosome 22. Flanking BanI restriction site, two arrows represent qPCR primer binding sites. Middle panel is the quantification of resection assay in the absence and presence of DNA DSB (-or + 4-OHT respectively). Values from siCTRL-3 + 4-OHT were considered to be 100%. Immunoblot controlling gene down regulation in DIvA cells transfected with siCTRL-3, siNLRP3-sp, siNLRP3-3 and siCtIP (right panel). ACTIN was used as a loading control. (A-B) shRNA-mediated NLRP3 knockdown was performed in MDA-MB-231. Mean and SEM of 2 (B) of 3 (A,C) biological replicates. (A-C) Mann Whitney test was performed. ns: nonsignificant, ** p < 0.01, *** p < 0.001, **** p < 0.0001.

It was recently reported that NLRP3 regulates part of the activity of ATM^19^. Since ATM inhibition is known to decrease RPA2 foci formation^27^, we examined whether the impact of NLRP3 on RPA2 recruitment following IR was due to its effect on ATM. As previously described, the ATM inhibitor (ATMi KU55933) impaired RPA2 recruitment to DSBs^27^, and this effect was even stronger in the absence of NLRP3, albeit at residual ATM activity^19^ (Figure 2B and Sup Figure 2F). We also confirmed the impact of ATMi on RAD51 recruitment compared with NLRP3 depletion. We similarly observed that the absence of NLRP3 dramatically reduced RAD51 foci formation compared with ATMi and the combination of both synergize, suggesting a possible role for NLRP3 independent of ATM regulation (Sup Figure 2E).

To determine whether this phenotype was due to reduced DNA end resection, we used a resection assay jointly developed by T. Paull and G. Legube on DIvA cells^28^. The amount of ssDNA decreased in the absence of CtIP used as a positive control for resection^29^. NLRP3 depletion also significantly decreased the amount of resection (Figure 2C). Altogether these results suggest that NLRP3 positively regulates the HR pathway by facilitating DNA end resection.

### NLRP3 associates with TRIP13 and REV7, two known actors of DSB repair

To further elucidate the function of NLRP3 in DDR, we sought NLRP3-binding partners by screening proteins that bind to FLAG-tagged NLRP3 (FLAG-NLRP3) in HeLa cells by mass spectrometry (Figure 3A). We identified TRIP13 among a group of proteins including the kinase NEK7, IPO5 and SUGT1 previously characterized as NLRP3-interactants^19,30–33^, validating our findings. Recently, the AAA+ ATPase remodeler TRIP13 has been reported to function in the DSB repair pathways via its control of REV7 conformation^14,15^. TRIP13 acts on HORMA family proteins, such as REV7 or MAD2^34^ by unwinding them to promote their conformational transition from closed to open, thereby inactivating them. Immunoprecipitation (IP) of FLAG-NLRP3 confirmed its interaction with endogenous TRIP13 (Figure 3B). We then performed a reciprocal IP by co-transfecting vectors expressing HA-tagged TRIP13 (HA-TRIP13) with FLAG-NLRP3 and showed that TRIP13 interacted with NLRP3 (Figure 3C).

**Figure 3.**
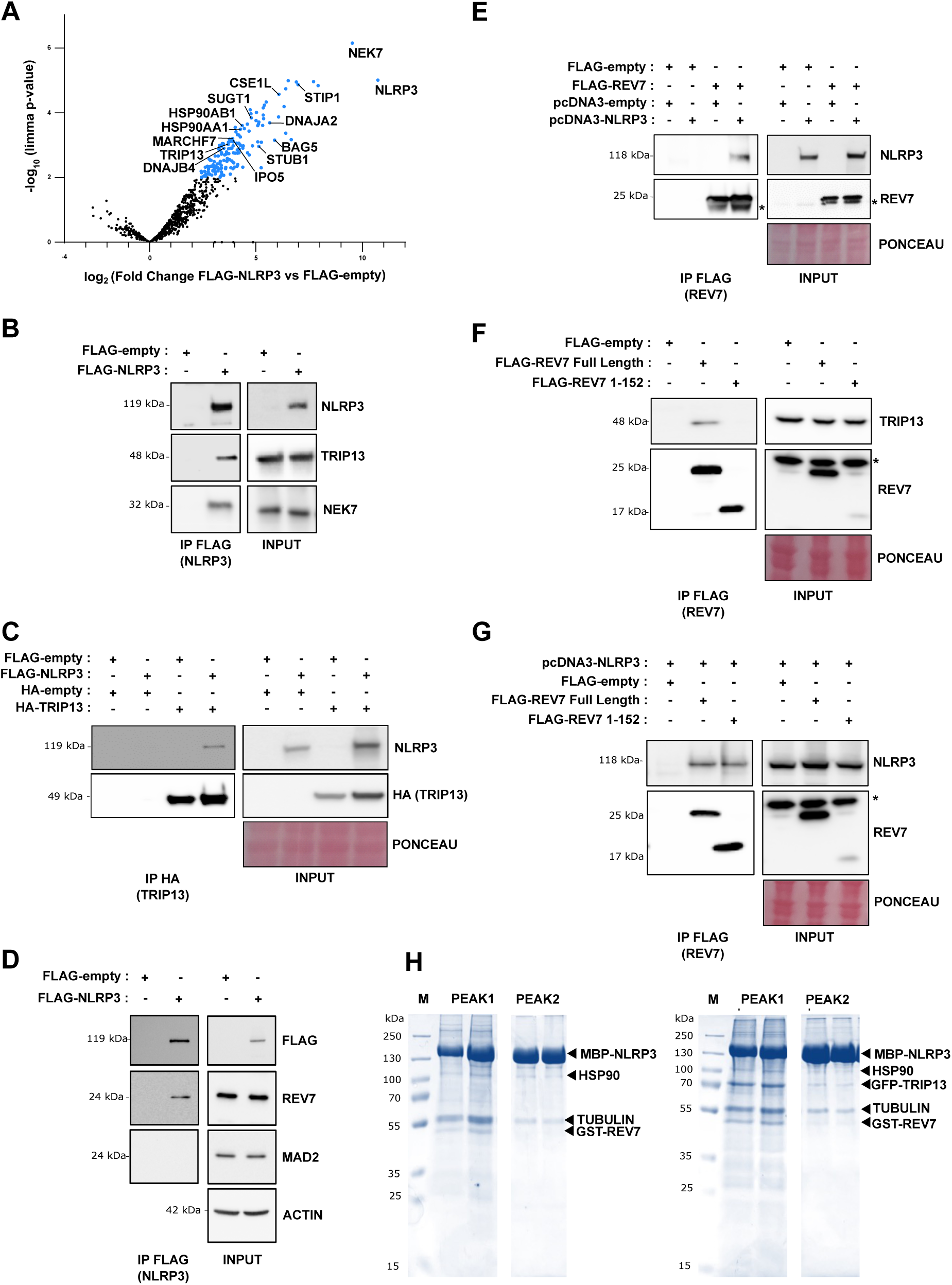
– NLRP3 interacts with TRIP13, and REV7. **(A)** MS-based proteomic identification of NLRP3 binding partners in HeLa cells. Volcano plot displaying the differential abundance of proteins in FLAG-NLRP3 and FLAG-empty (WT) co-immunoprecipitation eluates analyzed by MS-based quantitative proteomics. The volcano plot represents the −log10 (limma p-value) on y axis plotted against the log2 (Fold Change FLAG-NLRP3 versus FLAG-empty) on x axis. Blue dots represent proteins more abundant in FLAG-NLRP3 eluates (Fold change ≥ 4 and p-value ≤ 0.01, leading to a Benjamini-Hochberg FDR < 2%). n=3 independent experiments. **(B and D)** FLAG IP in HeLa cells transfected with FLAG-NLRP3. Eluates were analyzed by immunoblot with indicated antibodies. Representative of at least n=3 independent experiments. **(C)** Immunoblots of HA IP in HeLa cells transfected with FLAG-NLRP3 and HA-TRIP13. Representative of at least n=3 independent experiments. **(E)** Immunoblots of FLAG IP in HeLa cells transfected with untagged NLRP3 (pcDNA3-NLRP3) and FLAG-REV7. Representative of n=3 independent experiments. **(F)** Immunoblot of FLAG IP in HeLa cells transfected with FLAG-REV7 full length or with FLAG-REV7 1-152 aa (lacking the seatbelt region). Eluates were analyzed by immunoblot with indicated antibodies. Representative of n=2 independent experiments. **(G)** Immunoblots of FLAG IP in HeLa cells transfected with untagged-NLRP3 (pcDNA3-NLRP3) and FLAG-REV7 or FLAG-REV7 1-152 aa. Representative of two biological replicates. (E-G) * indicates non-specific band. **(H)** SDS-PAGE analysis of the two elution peaks from the dual MBP-NLRP3, GST-REV7 coexpression (left) or the triple MBP-NLRP3, GST-REV7, GFP-TRIP13 coexpression (right). Protein bands identified by mass spectrometry are indicated.

Since TRIP13 acts in DSB repair via its binding to REV7, we tested whether REV7 co-immunoprecipitated with FLAG-NLRP3, and confirmed that NLRP3 pulled down endogenous REV7 but not MAD2 (Figure 3D). We then used FLAG-tagged REV7 and immunoprecipitated NLRP3 (Figure 3E). To further explore these interactions, we generated a C-terminal deletion mutant of REV7 lacking the seatbelt region (REV7 1-152). As suggested in the literature, immunoprecipitation of REV7 1-152 did not pull-down TRIP13^35^, while it did pull-down NLRP3 implying that NLRP3 and TRIP13 interact with REV7 through different motifs (Figure 3F-G).

To determine whether the NLRP3 interaction with TRIP13 and REV7 is direct, we co-expressed MBP-NLRP3 and GST-REV7 or the three proteins MBP-NLRP3, GST-REV7 and GFP-TRIP13 in insect cells and purified the proteins via MBP affinity chromatography followed by size exclusion chromatography (SEC). Purified NLRP3 elutes in two peaks from SEC, peak 2 containing NLRP3 decamers involved in inflammasome assembly, while peak 1 contains more heterogeneous NLRP3 assemblies that exhibit ATP-hydrolysis activity^28^. Using Coomassie-stained SDS PAGE analysis we find that REV7 coeluted with NLRP3 species from peak 1, and to a lesser extend from peak 2, in the absence and in the presence of TRIP13, suggesting that NLRP3 does not require TRIP13 for binding to REV7 (Figure 3H). The protein identity was confirmed by peptide mass fingerprint analysis which also identified known cofactors HSP90α/β and α/β-Tubulin. In addition, we purified recombinant MBP-NLRP3 peak 1, GST-REV7 and GST proteins separately, incubated both protein pairs and performed MBP pull-down experiments using amylose beads. After three washing steps, a faint band of GST-REV7, but not the control GST, comigrated with MBP-NLRP3 indicating the direct binary interaction of the two proteins (Sup Figure 3A).

### NLRP3 and TRIP13 display distinct functions on HR

Given that TRIP13 loss was previously reported to either sensitize cells^15^ or not^14^ to PARPi, we compared the response of cells depleted in TRIP13 or in NLRP3 to control MDA-MB-231 cells using two PARPi. We observed no effect of TRIP13 depletion on olaparib sensitivity and a little effect on talazoparib sensitivity, compared with NLRP3 depletion (Sup Figure 4A, B). Consistent with these observations, TRIP13 depletion did not significantly decrease RPA2 foci formation compared with shNLRP3 in response to IR (Figure 4A and Sup Figure 4C). Hence, even though NLRP3 and TRIP13 both interact with REV7, they act differently on the HR pathway.

**Figure 4.**
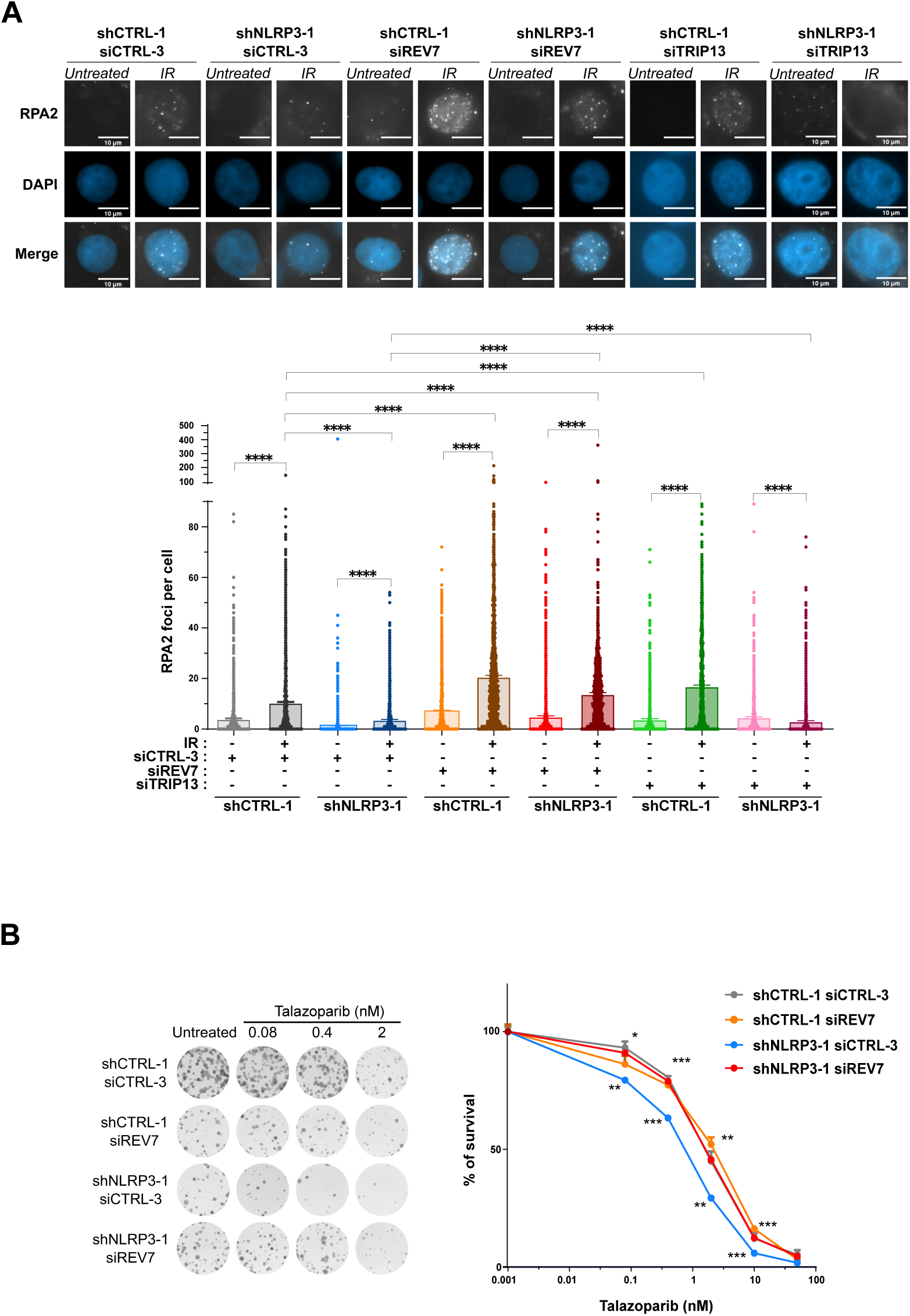
– Loss of REV7 restores homologous recombination in NLRP3-depleted cells. **(A)** Representative images of RPA2 foci 6 h after IR treatment (5 Gy) in shCTRL-1 and shNLRP3-1 MDA-MB-231 cells transfected with siCTRL-3, siTRIP13 or siREV7 (upper panel). DAPI was used to stain the nuclei. Scale bar, 10 μM. Quantification of RPA2 foci per nucleus is shown (lower panel). At least 100 cells were quantified per condition. **(B)** Sensitivity of shCTRL-1 and shNLRP3-1 MDA-MB-231 cells transfected with siCTRL-3 or siREV7 and treated with indicated doses of talazoparib using colony formation assay (CFA) (right panel). Representative images of clones are shown (left panel). (A-B) shRNA-mediated NLRP3 knockdown was performed in MDA-MB-231. Mean and SEM for at least three independent replicas is shown. The p-value correspond to a Mann Whitney’s test. ns: nonsignificant, ** p < 0.01, **** p < 0.0001.

### Loss of REV7 restores HR in NLRP3-depleted cells

One mechanism underlying PARPi resistance is HR restoration^36^. Given that REV7 depletion rescues HR in *BRCA1*-deficient cells^6^, we wondered whether this was similar in NLRP3-depleted cells and measured the formation of RPA2 foci in cells co-depleted for NLRP3 and REV7. Upon IR treatment, in shCTRL cells treated with siREV7, we first observed an increase in RPA2 foci formation. Importantly, REV7 depletion in NLRP3-deficient cells restored the defect observed on RPA2 foci formation at levels similar to the control condition (Figure 4A et Sup Figure 4C). To confirm this result, we tested the sensitivity of cells co-depleted for REV7 and NLRP3 to PARPi. We found that REV7 knockdown in shNLRP3 cells also abolished sensitivity to both talazoparib and olaparib (Figure 4B and Sup Figure 4D). Collectively these data show a functional interaction between NLRP3 and REV7, and support the notion that NLRP3 is a novel regulator of HR.

### NLRP3 inhibits REV7 recruitment to DSBs

Finally, we hypothesized that NLRP3 binding to REV7 may antagonize the shieldin complex. To determine whether NLRP3 affects the recruitment of REV7 to DSBs, we quantified REV7 foci after IR treatment, as previously demonstrated^15^. Interestingly, NLRP3 depletion resulted in increased REV7 foci formation after IR treatment (Figure 5A). As a control, cells were treated with NU7441, a DNA-PKcs inhibitor^37^, which decreased REV7 foci formation in both shCTRL and shNLRP3 conditions (Sup Figure 5A). The increase in REV7 foci in shNLRP3 was not caused by increased REV7 expression (Sup Figure 4C and Sup Figure 5B and 5C). To further establish the role of NLRP3 on REV7 recruitment to DSBs, we performed rescue experiments by overexpressing FLAG-NLRP3 in shNLRP3 cells. The addition of exogenous NLRP3 lowered the number of REV7 foci back to levels similar to those found in control conditions, demonstrating that NLRP3 controls REV7 recruitment to DSBs (Figure 5B and Sup Figure 5D).

**Figure 5.**
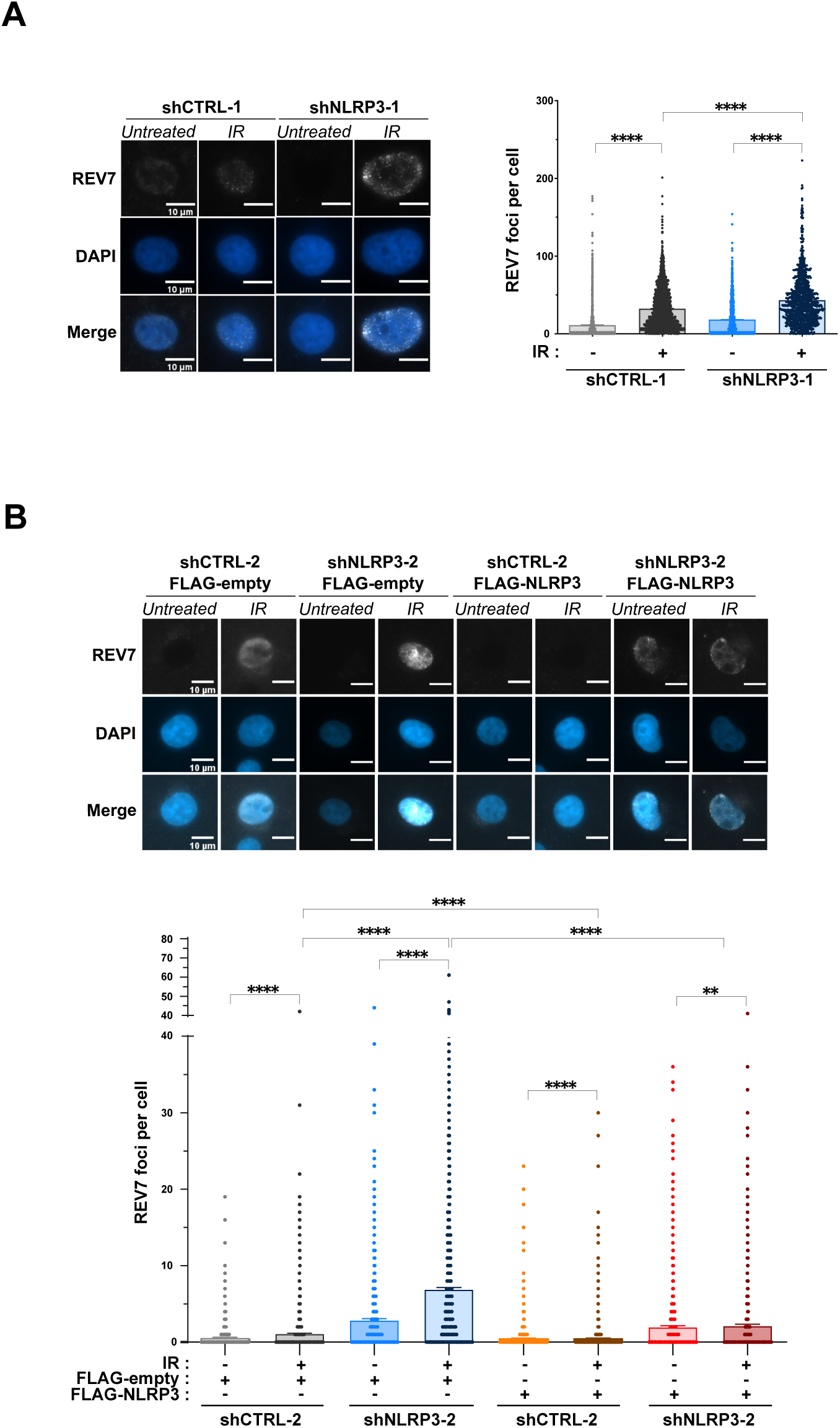
–NLRP3 inhibits REV7 recruitment to DSBs. (A) shCTRL-1 or shNLRP3-1 MDA-MB-231 cells were treated with 5 Gy IR and fixed at 6 h post-treatment. Representative images of REV7 foci (left panel). DAPI was used to stain the nuclei. Scale bar, 10 μM. Quantification of REV7 foci is shown (right panel). (B) shCTRL-2 or shNLRP3-2 MDA-MB-231 cells were transfected with a control or FLAG-NLRP3-expressing vector then treated with IR. Representative images of REV7 foci 6 h after IR (5 Gy) treatment (upper panel). DAPI was used to stain the nuclei. Scale bar, 10 μM. Quantification of REV7 foci is shown (lower panel). (A-B) Total number of foci are shown. At least 100 cells were quantified per experiment. Mean and SEM of n=3 (A) or n=2 (B) independent experiments. The p-value correspond to a Mann Whitney’s test. ** p < 0.01, **** p < 0.0001.

## DISCUSSION

The role of NLRP3 in inflammatory pathologies such as type 2 diabetes or Alzheimer’s disease has been well-established^18^. In the context of cancer, its role has mainly been addressed in the immune compartment^38^. Recent studies reported that NLRP3 is frequently silenced in solid tumors, including breast cancer, but its role in this context remains poorly understood^19,39^. Moreover, new functions in genome stability and DNA repair pathways are emerging. Recently, NLRP3 and NLRC4, another inflammasome sensor, were shown to contribute to ATM and ATR activation respectively in response to genotoxic stress in an inflammasome-independent way^19,40^. Here, we discovered that NLRP3 is located at the vicinity of DSBs and is part of the machinery involved in selecting the DSB repair pathway to promote the HR pathway. This new function is independent of the inflammasome complex. It is also independent of its role on ATM activation as ATMi only moderately altered RPA and RAD51 foci formation compared to the strong phenotype obtained after NLRP3 knockdown, which only dampens a fraction of ATM activity^19,40^ . Moreover, though REV7 recruitment was shown to rely partly on ATM activity, the increased REV7 foci formation in the absence of NLRP3 rules out a role via ATM^6^. In the literature, PRRs such as cGAS or AIM2 have been reported to play a role in DNA repair pathways. However, both receptors were shown to inhibit HR^41,42^. Hence, the role of NLRP3 in facilitating DNA end resection in the HR pathway and thus in maintaining global genome integrity is original.

Akin to TRIP13, NLRP3 is an AAA+ ATPase protein. Although we initially found an interaction with TRIP13, our results did not reveal a major role for TRIP13 in HR, as reported by de Krijger et al.^14,15^. However, we discovered that REV7 interacts with NLRP3 independently of TRIP13 and reported a functional interaction between both proteins. NLRP3 depletion increased the number of REV7 foci upon DSBs, while reducing DNA resection, suggesting that NLRP3 is a novel repressor of the shieldin complex.

Our findings are consistent with previous studies, which reported loss of REV7 or shieldin subunits as one of the mechanisms allowing HR restoration in a *BRCA1*-null background^6,9^. We showed that NLRP3 co-depletion with REV7 restores resection as assessed by RPA2 foci and PARPi resistance. The precise mechanism used by NLRP3 to impair the function of REV7, however, remains to be evaluated. We propose that its interaction with REV7 prevents the assembly of the shieldin complex. Finally, since REV7 forms different protein complexes, whether NLRP3 controls other REV7 functions such as its role in the DNA polymerase zeta complex in the translesion DNA synthesis^15,43^ is an important question for future research. While many mechanisms leading to HRD and resistance to PARPi have been identified, the discovery of novel therapeutic targets such as NLRP3 offers new and unexplored perspectives. Further investigations into NLRP3 inhibition to improve the efficacy of PARPi could be of major importance.

## MATERIALS AND METHODS

### Cell cultures

HeLa (human mammary adenocarcinoma cells, U-2 OS (human osteosarcoma cells), 293T (Human embryonic kidney 293T) and THP-1 (monocyte) were purchased from American Type Culture Collection (ATCC). HMEC-hTERT (human mammary epithelial cells) and MDA-MB-231 were obtained from Alain Puisieux’s laboratory^44,45^. DIvA cells were obtained from Gaëlle Legube’s laboratory.

MDA-MB-231, HeLa, U-2 OS and 293T were grown in Dulbecco’s modified Eagle medium (DMEM) (Gibco, catalog no.41965-039), supplemented with 10% FCS (Gibco, catalog no.10270-106), 1% Glutamax (Gibco, catalog no.35050-038), 1% Penicillin-Streptomycin (Gibco, catalog no.15140-122) and 1% sodium pyruvate (Gibco, catalog no.11360-039). HMEC-hTERT were cultured in DMEM/Ham F12 medium (Gibco, catalog no.10565018), supplemented with 1% GlutaMAX, 10 ng/mL human EGF (PromoCell), 0.5 mg/mL hydrocortisone (Sigma-Aldrich, catalog no. H0888), and 10 mg/mL insulin (Actrapid). THP-1 were cultured as previously described^46^. Cell lines were grown adherently at 37°C in a humid atmosphere and 5% CO_2_. Mycoplasma contaminations were tested monthly (Lonza, catalog no.LT07-318).

### Lentiviral production

293T cells at 70% confluency were transfected with 5 μg plasmid DNA in 10 cm dish using polyethyleneimine (PEI) (Tebu-bio, catalog no.23966-2) and lentiviral packaging plasmid mix (second-generation packaging system psPAX and pMD2.G plasmids) (Table 1). Supernatant containing viral particles was collected 48 h post transfection, filtered twice through 0.45 µm filters, supplemented with 6 µg/µL polybrene (Sigma, catalog no TR-1003-G) and added to MDA-MB-231 and U-2 OS cells. Cells were transduced with shRNA lentiviruses targeting human NLRP3 or a scrambled control hairpin. Cells were selected with 10 μg/mL of blasticidin (InvivoGen, catalog no.ant-bl-1) or 1 μg/mL of puromycin (InvivoGen, catalog no.ant-pr-1).

**Table 1.**
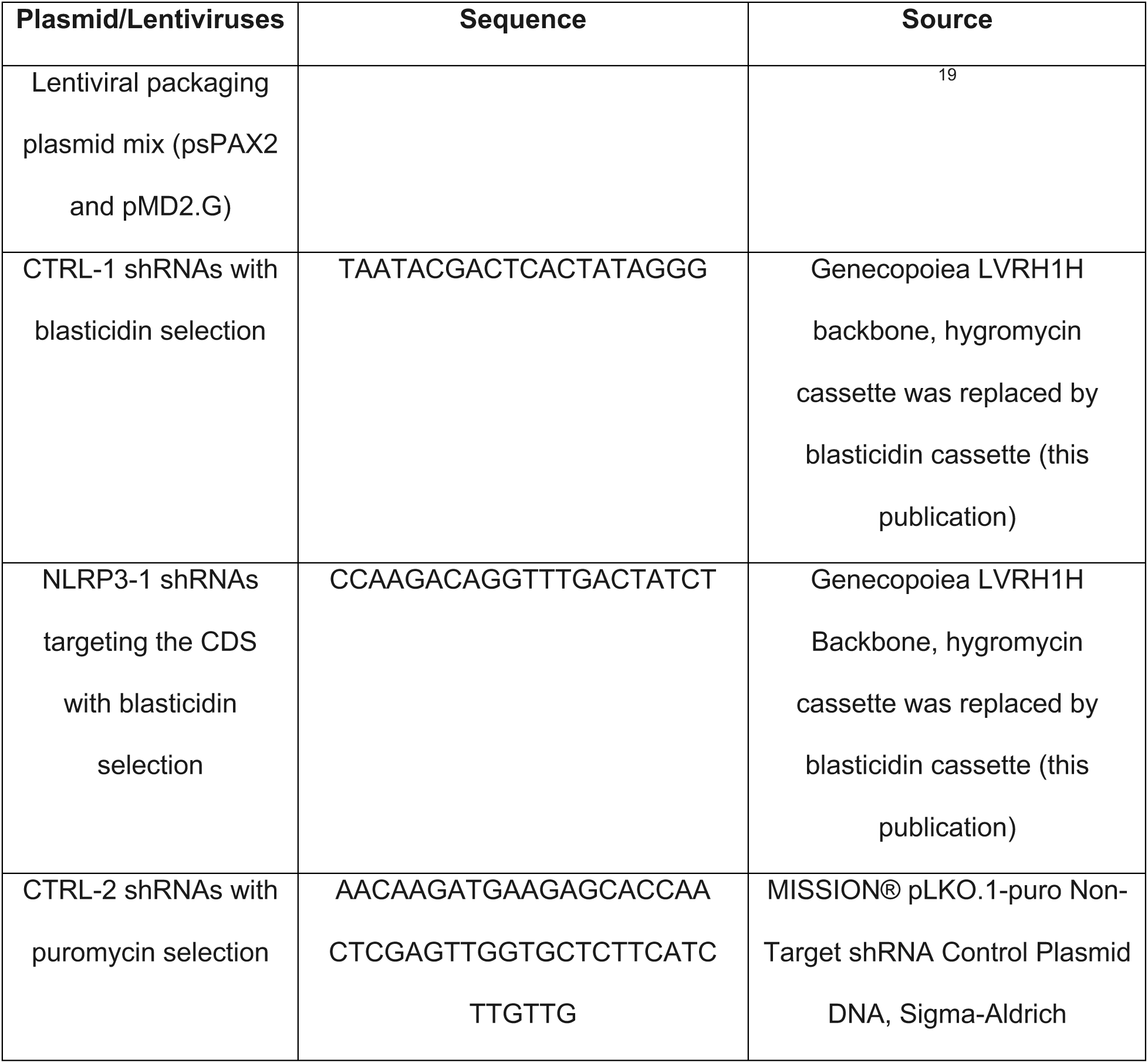
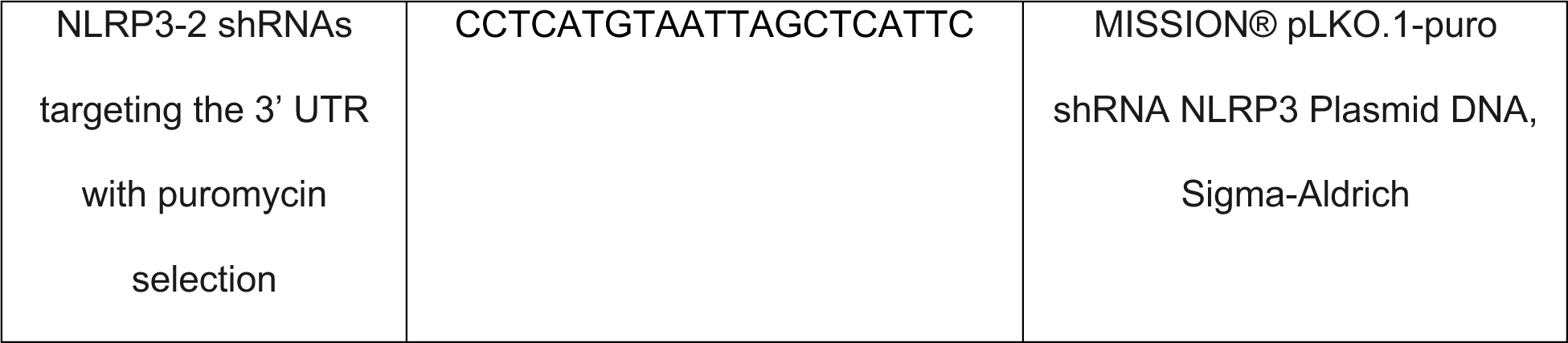

### Cell Transfection siRNA

4×10^5^ and 2×10^5^ HeLa/U-2OS and MDA-MB-231 cells were seeded per well, respectively, in 6-well plates, approximately 60% confluence. Cells were transfected with specific siRNA duplex at a concentration of 20 nM (Table 2). The transfection was performed in adherent cells with INTERFERin (Ozyme, catalog no.101000016) according to the manufacturer’s instructions.

**Table 2.**
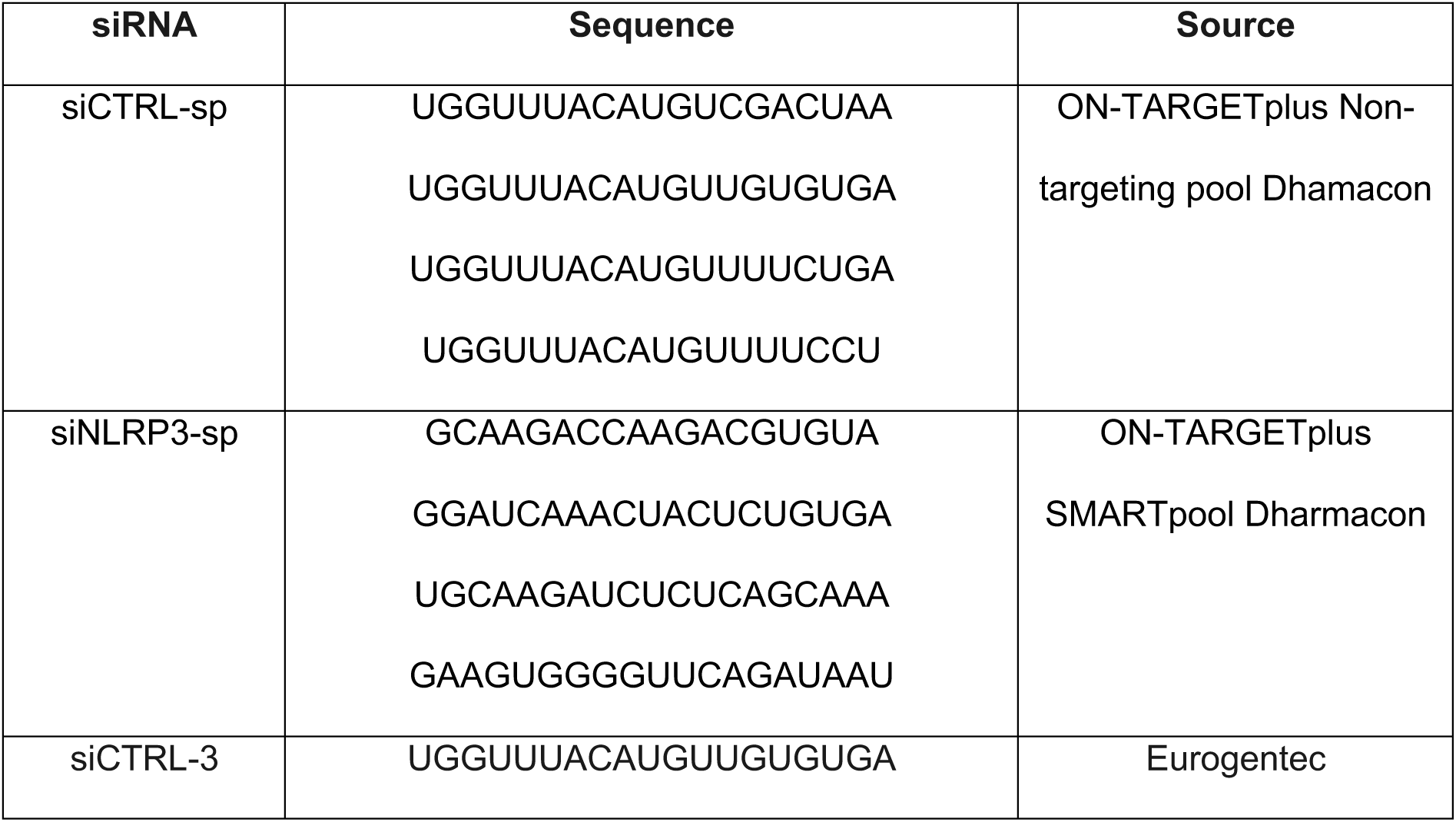
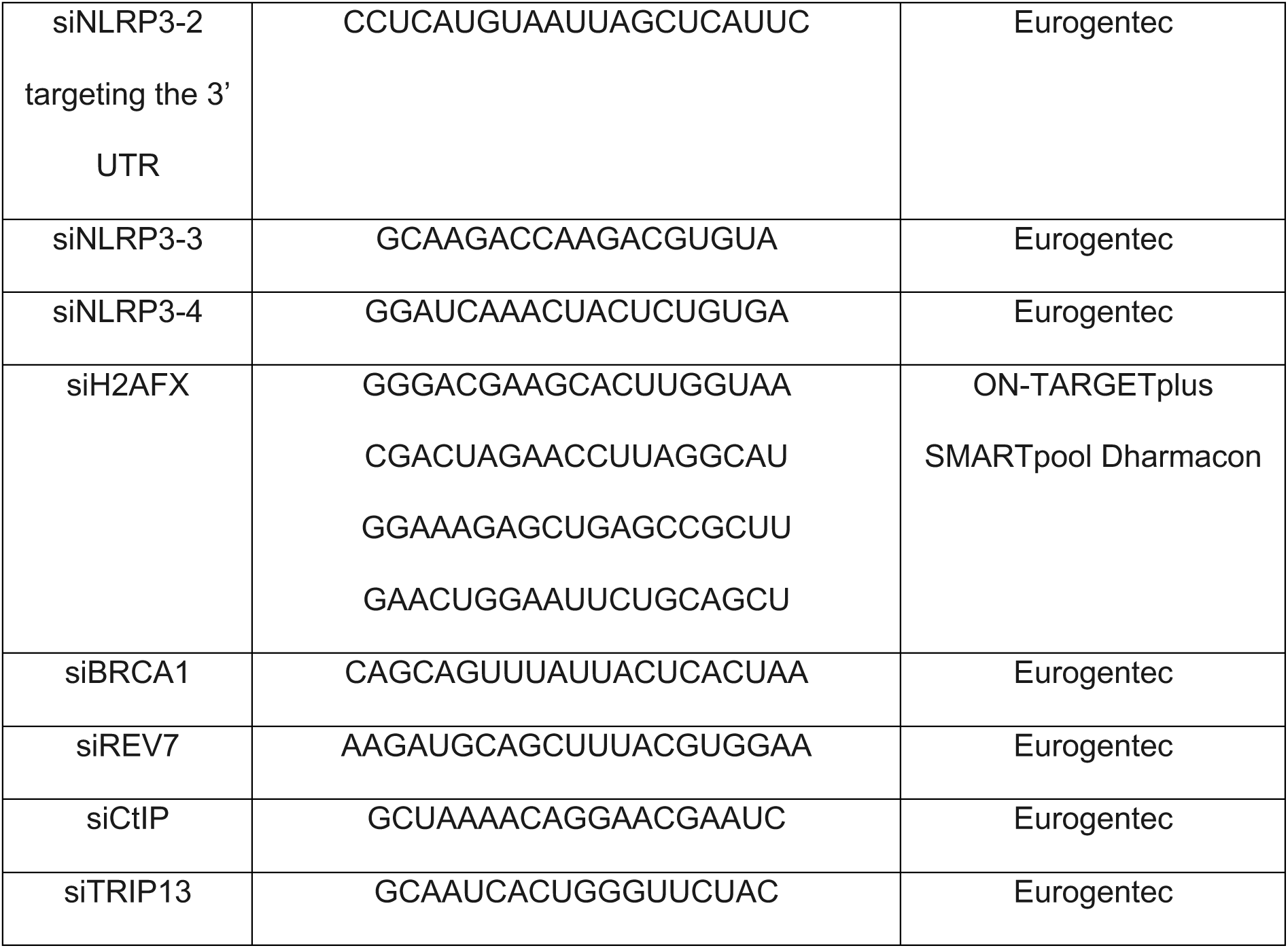

### Plasmids

1.5×10^6^ HeLa and MDA-MB-231 cells were seeded per 10 cm dish. The cells were transfected with a plasmid using with Lipofectamine 2000 (Invitrogen, catalog no.11668019) as per manufacturer’s instructions.

GFP-REV7 was purchased from Addgene (#114128 pDEST-FRT/TO-eGFP-REV7). From this vector, the REV7 cassette was amplified and the FLAG full length or FLAG REV7 Δ seat belt motif (REV7 1-152) was cloned into pCI-Neo plasmids (Table 3). Since there was no immediate enzyme site upstream of the gene, the cloning strategy was adapted by cleaving a part of the vector and re-ligating it with the REV7 Δ seat belt motif fragment (REV7 1-152) and full-length. To do so, PCR amplification was conducted using the same vector as a template with forward 5’ TATCACGAGGCCCTTTCG 3’ and reverse 5’GCGGCCGGCCTACGTGTGCACCAGGACTGTGAA 3’ primers. The vector and PCR were then digested by AscI and NaeI enzymes and ligated after purification. Positive clones were selected and confirmed by restriction digestion. Finally, the sequence was confirmed by sequencing the cloned regions.

**Table 3.**

| Plasmid | Source |
| --- | --- |
| pCR3-FLAG-NLRP3 | 19 |
| pCR3-FLAG-empty | 19 |
| pcDNA3-NLRP3 | Gift from Bénédicte Py laboratory |
| pcDNA3-empty | Gift from Kyle Miller laboratory |
| pRP-HA-TRIP13 | This publication: Vector Builder pRP-Bsd-CMV-HA/hTRIP13 (nm_004237,4) |
| pcDNA3-mCherry-empty | 19 |
| pcDNA3-mCherry-NLRP3 | 19 |
| pcDNA3-HA-empty | Gift from Katja Lamia Laboratory |
| pDEST-GFP-REV7 | Addgene pDEST-FRT/TO-eGFP-REV7 #114128 |
| pCI-FLAG-REV7 | This publication |
| pCI-FLAG-REV7ΔSeatbelt (REV7 1-152) | This publication |

To generate the shNLRP3-2 plasmid with blasticidin resistance, the hygromycin cassette was removed from the commercial vector psi-LVRH1H by digesting it with *AgeI* and *MluI* enzymes. The blasticidin resistance cassette was amplified by PCR using the following primers F-5’ CCATACCGGTACCATGGCCAAGCCTTTGTCTCAA 3’, R-5’ CAGACGCGTTTAGCCCTCCCACACATAACCAGA 3’ from pLKO.1-Blasticidin vector (Addgene #26655), and ligated after *AgeI* and *MluI* digestion and gel purification with the vector backbones. Positive clones were checked by double digestion and confirmed by sequencing.

### Drugs

48 h after transfection, cells were treated with etoposide (TEVA santé 100 mg/5 mL) at a concentration of 10 µM or 100 µM. Before irradiation, cells were treated for 1 h with a specific ATM inhibitor (KU55933, Selleckchem, catalog no.S1092) or DNA-PKcs inhibitor (NU7441/KU-57788, Selleckchem, catalog no.S2638) at a concentration of 10 μM and 1 μM, respectively.

Cells were treated during 16 h before irradiation with MCC950 (Selleckchem, catalog no.S8930), a selective inhibitor of NLRP3 inflammasome activation at a concentration of 1μM^26^.

For clonogenic survival assays, cells were transfected with siRNA. 24 h later, cells were seeded at low density (200 cells per well into 6-well). 48 h after transfection, cells were treated with olaparib (AZD2281 Selleckchem, catalog no.S1060) at concentrations ranging from 0.0001 µM to 10 μM or with talazoparib (BMN 673, Selleckchem, catalog no.S7048) at concentrations ranging from 0.001 nM to 100 nM.

### Irradiations

Cells were irradiated using a 6-MeV γ-ray clinical irradiator (SL 15 Phillips) at the Centre Léon Bérard, with a dose rate of 6 Gy.min-1 to obtain the required dose.

### Proximity Ligation Assay

Cells were grown on glass coverslips in 6-well plates, then washed twice in PBS and fixed in PBS-Paraformaldehyde 4% for 15 min. Cells were washed twice with PBS before being permeated for 3 min with Triton 0.5% in PBS. The manufacturer protocole was adapted as follow. Cells were incubated with Duolink® blocking solution (1 drop per 1 cm^2^) for 1 h at 37°C in a humidified chamber. They were then incubated with primary antibodies against NLRP3 (Millipore, catalog no.ABF23, 1:500) and γH2AX (Sigma, catalog no.05-636, 1:500) in Duolink® antibody diluents for 1 h at 37°C in a humidified chamber. After three washes in Buffer A, the affinity purified goat anti-rabbit IgG Duolink in Situ PLA probe PLUS® and affinity purified goat anti-mouse IgG Duolink in Situ PLA probe MINUS® (1:5), containing the secondary antibody conjugated with complementary oligonucleotides, were added and incubated for 1 h at 37°C in a humidified chamber. Subsequently, cells were washed three times in Buffer A, and incubated in 1 unit/μL of T4 DNA ligase in diluted ligase buffer (1:5) for 30 min at 37°C in a humidified chamber. After three washes in Buffer A, cells were incubated in 5 units/μL of DNA polymerase in diluted polymerase buffer with red fluorescent-labeled oligonucleotides (1:5) for 100 min at 37°C. A final round of washes was performed twice for 10 min in Buffer B, then 1 min in 0.01x Buffer B. The slides were then mounted using 5 μL of Duolink in situ mounting medium containing DAPI. Nail polish was used to seal the edges of the coverslips. All of the reagents are included in the PLA kit (Sigma, DUO92106, DUO92004, DUO92002, DUO92008, DUO82049, DUO082047, DUO82040)^47^.

Imaging of the slides was performed using a Nikon Eclipse Ni-E or Zeiss Axio Imager M2 microscope. Images were acquired under identical conditions at x60 magnification. At least 100 cells were analyzed per condition. The analysis of the number of dots per cell was performed using the Image J software. Quantification was done using the macro published by Poulard et al^47^. Data analyses were performed using Prism Version 10.0 (GraphPad Software).

### Clonogenic survival assays

MDA-MB-231 or U-2 OS cells were seeded in 10 cm dishes and then transfected with siRNA as indicated. 48 h after the transfection, cells were trypsinized, counted and seeded at low density; 200-2,000 cells/ 6-well plates (MDA-MB-231) or 600-1,200 cells/6-well plates (U-2 OS). The day after seeding, cells were treated with either DMSO or the PARP inhibitor (olaparib or talazoparib) at the indicated concentrations. Each experiment was performed at least 3 times with 6 wells per condition. Two weeks after, cells were harvested, fixed and stained for 1 h at room temperature with solution containing 50% ethanol, 5% acetic acid, and 0.5% brilliant blue R (Sigma-Aldrich; catalog no. B7920), washed in water, and air-dried overnight. Clone counting was done manually and the percentage survival took into account the number of clones seeded as well as the plating efficiency. Analysis was carried out on Prism Version 10.0.

### DSB repair assays

For HR repair assays, U-2 OS DR-GFP cells were described previously^23^. These cells were transfected with mCherry-NLRP3 or the mCherry-control vector as indicated along with either HA I-SceI or empty vector^48^ by using PEI or with indicated siRNA prior to HA I-SceI or control vector transfections. 24 h after transfection, cells were trypsinized and half of the cells were harvested to verify HA I-SceI expression or co-expression with mCherry. The other half of the cells were re-plated and harvested after 72 h of transfection to measure GFP-expressing cells. For HA detection by flow cytometry, rat anti-HA primary (Roche #21319000) and goat anti-rat AlexaFlour-647 (Invitrogen #A21247) secondary antibodies were used. The percentage of HR repair post-I-SceI cleavage was calculated by dividing the percentage of GFP+-cells 72 h by the percentage of HA-positive and mCherry-positive cells at 24 h. Protein expression or depletion were confirmed by immunoblotting. Results were normalized against HA I-SceI treated mCherry control vector or siCTRL.

### MS-based quantitative proteomic analyses

Three independent co-immunoprecipitation experiments from HeLa cells expressing FLAG-NLRP3 or FLAG alone were performed. The eluted proteins, solubilized in Laemmli buffer, were stacked in the top of a 4-12% NuPAGE gel (Invitrogen). After staining with R-250 Coomassie Blue (Biorad), proteins were digested in-gel using trypsin (modified, sequencing purity, Promega), as previously described^49^. The resulting peptides were analyzed by online nanoliquid chromatography coupled to MS/MS (Ultimate 3000 RSLCnano and Q-Exactive Plus, Thermo Fisher Scientific) using a 140 min gradient. For this purpose, the peptides were sampled on a precolumn (300 μm x 5 mm PepMap C18, Thermo Fisher Scientific) and separated in a 75 μm x 250 mm C18 column (PepMap, 3µm, Thermo Fisher Scientific). MS and MS/MS data were acquired by Xcalibur (Thermo Fisher Scientific).

Peptides and proteins were identified by Mascot (version 2.8, Matrix Science) through concomitant searches against the Uniprot database (Homo sapiens taxonomy, March 2024 version) and a homemade database containing the sequences of classical contaminant proteins found in proteomic analyses (human keratins, trypsin, etc.). Trypsin/P was chosen as the enzyme and two missed cleavages were allowed. Precursor and fragment mass error tolerances were set at 10 and 20 ppm, respectively. Peptide modifications allowed during the search were: Carbamidomethyl (C, fixed), Acetyl (Protein N-term, variable) and Oxidation (M, variable). The Proline software version 2.2.0^50^ was used for compiling, grouping, and filtering the results (conservation of rank 1 peptides, peptide length ≥ 6 amino acids, false discovery rate of peptide-spectrum-match identifications < 1%^51^, and minimum of one specific peptide per protein group). Proline was then used to perform a MS1 label-free quantification of the identified protein groups based on razor and specific peptides.

Statistical analysis was then performed using the Prostar software version 1.30.5^52^. Proteins identified in the contaminant database, proteins identified by MS/MS in less than two replicates of one condition, and proteins detected in less than three replicates of one condition were discarded. After Log2 transformation, abundance values were normalized by condition using median quantile before missing value imputation (slsa algorithm for partially observed values in the condition and DetQuantile algorithm for totally absent values in the condition). Statistical testing was then conducted using Limma, whereby differentially expressed proteins were sorted out using a Log2 (fold change) cut-off of 2 and a p-value cut-off of 0.01, leading to a false discovery rate (FDR) inferior to 2% according to the Benjamini-Hochberg estimator. Differentially abundant proteins identified by MS/MS in less than two replicates and detected in less than three replicates in which they were more abundant were manually invalidated (p-value = 1). The mass spectrometry proteomics data have been deposited to the ProteomeXchange Consortium via the PRIDE partner repository with the dataset identifier PXD058288.

### Immunoprecipitation

Anti-Flag M2 Affinity Gel beads (Sigma Aldrich, catalog no. A2220) or anti-HA-Agarose beads (Sigma Aldrich, catalog no.P3296) were washed twice with IP buffer (Tris HCl 100 mM pH 8.0, MgCL_2_ 10 mM, NaCl 90 mM, Triton X-100 0.1%)^19^ by centrifugation for 3 min, 5,000*g* at 4°C. 30 h post-transfection, cells were washed in PBS 1X, incubated in IP buffer and gently agitated at 4°C for 30 min. Cell extracts were collected and centrifuged 15 min, at 10,000*g* at 4°C to removed insoluble fractions. The supernatant was recovered, a small fraction was kept for input, and incubated with antibody-coupled beads overnight, on a rotating wheel, at 4°C. The beads were then washed three times in IP buffer on a rotating wheel for 15 min at 4°C, followed by centrifugation for 5 min at 5,000*g* at 4°C. IP proteins were denatured in 6X Laemmli buffer for 10 min at 95°C. Samples were analyzed using immunoblot as described below.

### Protein purification

Recombinant MBP-tagged hNLRP3 (amino acids 3-1036, UniProt accession number Q96P20) was expressed in *Sf9* insect cells with the Bac-to-Bac expression system for insect cell infection as described^53^. Baculovirus was amplified over two passages and then added to constitute 5% of the final insect cell culture. After 72 hours incubation at 27°C, cells were harvested by centrifugation at 2000 rpm for 20 min at 4°C.

For protein purification, the resulting pellet was resuspended in purification buffer (50 mM HEPES, 150 mM NaCl, 10 mM MgCl_2_, 1 mM ADP, 0.5 mM TCEP). After adding PMSF to a final concentration of 100 µM cells were disrupted by sonication. The lysate was centrifuged at 25,000 rpm for 1 h at 10°C. The supernatant was passed through a 0.22 µM filter and loaded on a 5 ml MBPtrap column. After washing out unbound protein, the MBP-hNLRP3 was eluted using 15 ml purification buffer supplemented with 15 mM maltose. The eluted protein was concentrated with a centrifugal filter to approximately 5 mg/ml and then subjected to further purification via gel filtration on a Superose 6 increase column.

The same expression and purification procedures were followed for GST-tagged hREV7 (amino acids 1-211, UniProt accession number Q9UI95) using a GSTtrap column for affinity purification. For coexpression of both proteins, second-passage baculovirus of both virus lines were combined and added to *Sf9* insect cells. Same applies the coexpression of both proteins together with GFP-TRIP13 (amino acids 1-432, UniProt accession number Q15645). The purification was conducted following the described protocol using a MBPtrap column. Stained protein bands were cut-out from SDS PAGE gels and subjected to bioanalytical peptide mass fingerprint spectrometry for the determination of the protein identity.

### Pull-down

To investigate the interaction between MBP-hNLRP3 and GST-hREV7 *in vitro*, 6 µM MBP-hNLRP3 was incubated with either 12 µM GST-hREV7 or GST as a control for 1 hour on ice. Subsequently, 50 µl amylose resin was added to these samples, followed by 1 hour incubation on ice. The beads were washed three times with 200 µL purification buffer, with samples collected after the first and third wash steps. For protein elution, 50 µl maltose supplemented purification buffer was added and samples were taken after 30 min. Samples were analyzed by SDS-PAGE followed by Coomassie staining.

### Immunofluorescence

2.5×10^5^ and 4×10^5^ MDA-MB-231 and U-2 OS cells per well were plated on glass coverslips in 6-well plates, respectively. They were then transfected with siRNA or/and pre-treated with drugs and either left untreated or treated at 5 Gy IR. After 1 h (for 53BP1) or 6 h (for RPA2 and REV7), they were washed twice with PBS and pre-extracted with 0.5% Triton X-100 (Sigma, catalog no.9036-19-5) (for RPA2 and REV7) in PBS for 5 min, followed by 4% paraformaldehyde (Electron Microscopy Sciences, catalog no.15710) fixation for 20 min at 4°C, and blocked with 3% BSA (Roche, catalog no.10735094001) in PBS for 1 h at room temperature (for RPA2, REV7 and RAD51) or 2% BSA with Tween 0.05% (for 53BP1) (Table 4). For RAD51 no pre-extraction was performed, cells were fixed after 6 h. Cells were lysed in a lysis buffer (Sucrose 300 mM, MgCl2 3 mM, Tris pH 7.0 20 mM, NaCl 50 mM, Triton X100 1%) followed by overnight incubation in primary antibody and blocking solution at 4°C, and in secondary antibody for 1 h at room temperature. Coverslips were mounted with mounting medium containing DAPI (Sigma, catalog #DUO82040) and sealed with nail polish (Electron microscopy sciences, catalog no.72180). Images were acquired using a Nikon Eclipse Ni-E or Zeiss Axio Imager M2 fluorescence microscope at x60 magnification. Foci were counted using ImageJ software. At least 50 cells were counted for each sample. Data analyses were performed using Prism Version 10.0 (GraphPad Software).

**Table 4.**

| **Antibodies** | **Clone** | **Identifier** | **Source** | **Dilution** |
| --- | --- | --- | --- | --- |
| Anti-RPA2 |  | MABE285 | Millipore | 1/500 |
| Anti MAD2L2/REV7 | EPR13657 | ab180579 | Abcam | 1/200 |
| Anti-RAD51 | PC130 | ab-1 | Calbiochem | 1/200 |
| Mouse anti human 53BP1 |  | MAB3802 | Millipore | 1/500 |
| Alexa Fluor 488 goat anti-rabbit IgG |  | A11034 | Invitrogen | 1/800 |
| Alexa Fluor 488 goat anti-mouse IgG |  | A11029 | Invitrogen | 1/800 |

### Q-PCR based end resection assay

DIvA (AsiSI-ER-U-2 OS) cells were seeded at a confluency of 30% one day before siRNA transfection. The next day siRNA-mediated gene depletion was performed using 20 nM final concentration of siRNA oligos and Interferin following the manufacturer’s protocol. After 24 h, cells were separated and grown on 2 plates. On day 3, a second siRNA transfection was performed with siRNA oligos at 10 nM. On day 4, one of the plates was treated with 0.5 µM 4-hydroxytamoxifen (ref), the other plate left as control. After 4 h, all plates were harvested using trypsin-EDTA, and gDNA were extracted using the Qiagen gDNA extraction kit (ref) following the manufacturer’s protocol except no vortexing was performed. Genomic DNA extraction was performed on 2×10^6^ cells. Before performing q-PCR, gDNA was treated with RNaseH by taking 3 µg of gDNA in 100 µL reaction volume and incubated for 1 h at 37°C, then heat denatured at 65°C for 20 min. 500 ng of RNase treated DNA was then used for AsisSI along with either BanI digestion or mock digestion. For q-PCR, 40 ng of DNA per well was taken in triplicate to amplify the specific region of genomic DNA and the results were compared to the control sample. To calculate the percentage of ssDNA generated for each sample, we applied the following the formula described in^28^ % ssDNA = 1/((POWER(2,Ct of BanI digested sample – Ct of non-digested sample – 1) + 0.5)) × 100. For normalization, tamoxifen-treated siCTRL cells were considered as 100 percent.

### Comet Assay

Alkaline comet assay was performed following the manufacturer’s instructions (Bio-techne, catalog no.4250-050-K). MDA-MB-231 cells were treated for 3 h with etoposide (100 μM) or irradiated 4 h (5 Gy) before being pelleted and resuspended in cold PBS at a concentration of 2×10^6^ cells/mL. Resuspended cells were then mixed with pre-warmed 1% low-melting point agarose. The mix was then spread on a warm slide pre-coated with agarose. After 30 min at 4°C, slides were immersed in cold lysis solution for 1 h at 4°C in a dark chamber. The slides were then denatured in an alkaline buffer (NaOH 0.3 M, EDTA 1 mM) for 1 h in the dark at 4°C. The slides were subjected to electrophoresis in alkaline buffer for 30 min at 25 volts/300 mA at 4°C. After migration, slides were immersed in neutralizing buffer (0.4 M Tris, pH 7.5) for 10 min at 4°C. Finally, the cells were stained with Sybr Gold (Invitrogen, catalog no.S11494). Images were observed using a Nikon Eclipse Ni-E fluorescence microscope at 40x and analysis was performed on at least 50 comets per condition with OpenComet on ImageJ software.

### Western blot analysis

Cells were washed with PBS and lysed directly on ice in Laemmli 2x buffer (0.125 M Tris HCl pH 6.8, 2% SDS, 0.5 M DTE). Proteins were quantified using the Bradford’s reagent (Biorad, catalog no.5000006, 1 :5) and a standard curve using quantified Bovine Serum Albumin (Sigma, catalog no.A9647). Optical density was read at 450 nm (with MultiskanTM FC Microplate photometer, ThermoFisher Scientific, catalog no.51119000). Prior to migration, 6x Laemmli Buffer (0.35 M Tris-HCl pH 6.8, 10% SDS, 50% glycerol, 0.5 M DTE, 0,004% bromophenol blue) was added to the samples and then denatured for 5 min at 95°C. Proteins were resolved using 8% (for the higher molecular weight proteins: NLRP3, CtIP, TRIP13) or 15% (for the lower molecular weight proteins: REV7, MAD2, NEK7, ASC, ACTIN, α-TUBULIN) SDS-PAGE gel (Table 5). Migration was performed in 1x Tris-glycine buffer (25 mM Tris-base, 192 mM glycine, 0.1% SDS). The proteins were then transferred to nitrocellulose membranes using Trans-Blot® Turbo (TM Transfer System, Biorad, catalog no.170-01918). Membranes were blocked using 1x TBST (0.2 mM Tris Base pH 7.5, 0.001% Tween, 0.5 M NaCl) 5% milk and incubated overnight with primary antibodies (Table 5) at 4°C, secondary. Horseradish peroxidase-coupled antibodies were from Promega (anti-mouse and anti-rabbit IgG HRP conjugate, catalog no. W402B and W401B respectively) and chemiluminescence reagent from Biorad (Clarity TM Western ECl Substrate, catalog no.170-5061). Visualization was performed on a ChemiDoc Imaging System (Biorad) or using Amersham Hyperfilm^TM^ ECL (GE Healthcare), and signal intensities were quantified using the Image Lab software (Biorad).

**Table 5.**
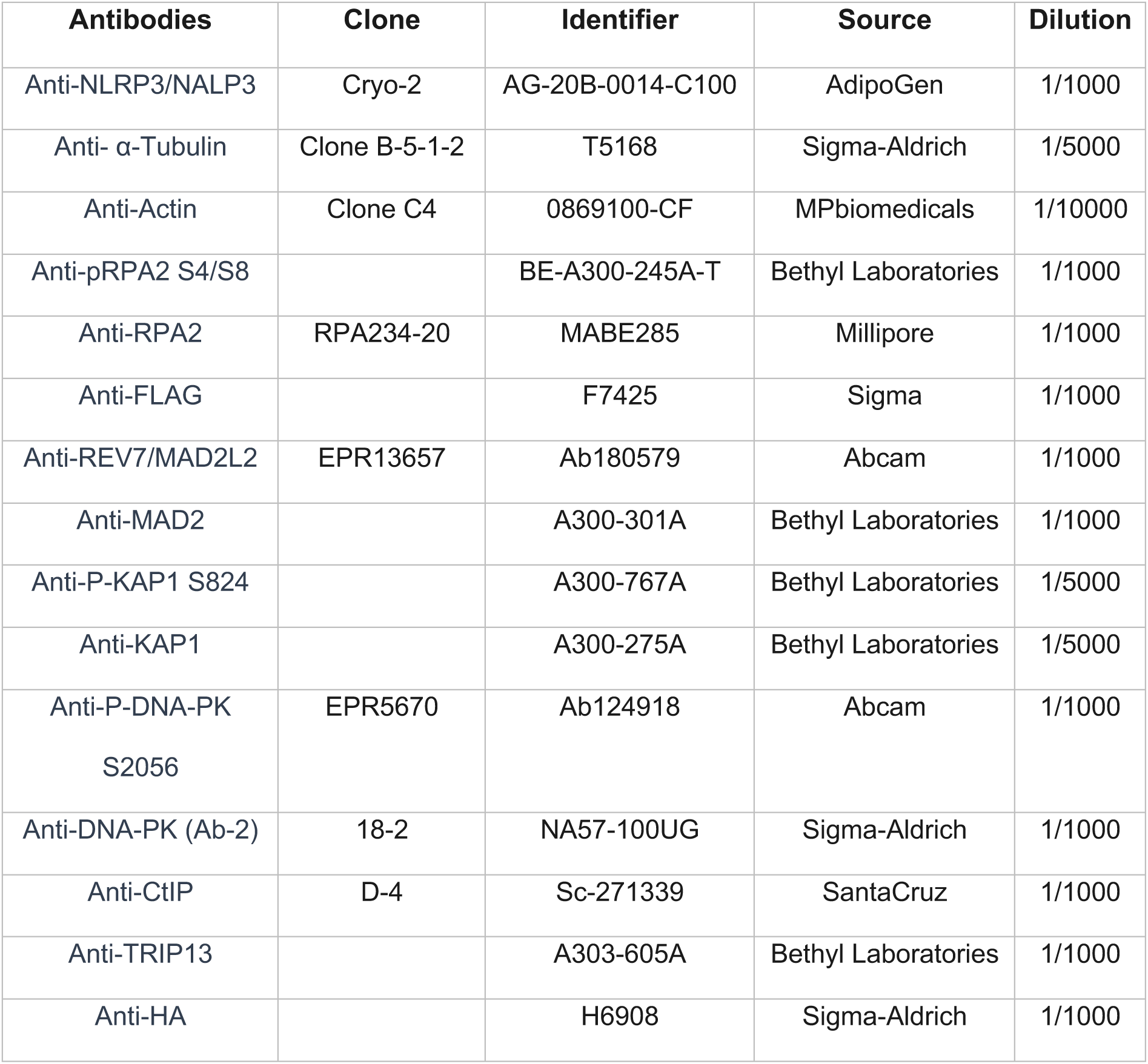
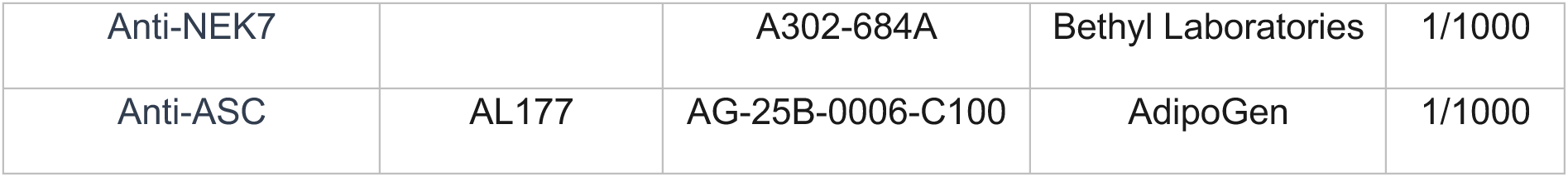

| **Antibodies** | **Clone** | **Identifier** | **Source** | **Dilution** |
| --- | --- | --- | --- | --- |
| Anti-NLRP3/NALP3 | Cryo-2 | AG-20B-0014-C100 | AdipoGen | 1/1000 |
| Anti- α-Tubulin | Clone B-5-1-2 | T5168 | Sigma-Aldrich | 1/5000 |
| Anti-Actin | Clone C4 | 0869100-CF | MPbiomedicals | 1/10000 |
| Anti-pRPA2 S4/S8 |  | BE-A300-245A-T | Bethyl Laboratories | 1/1000 |
| Anti-RPA2 | RPA234-20 | MABE285 | Millipore | 1/1000 |
| Anti-FLAG |  | F7425 | Sigma | 1/1000 |
| Anti-REV7/MAD2L2 | EPR13657 | Ab180579 | Abcam | 1/1000 |
| Anti-MAD2 |  | A300-301A | Bethyl Laboratories | 1/1000 |
| Anti-P-KAP1 S824 |  | A300-767A | Bethyl Laboratories | 1/5000 |
| Anti-KAP1 |  | A300-275A | Bethyl Laboratories | 1/5000 |
| Anti-P-DNA-PK<br>S2056 | EPR5670 | Ab124918 | Abcam | 1/1000 |
| Anti-DNA-PK (Ab-2) | 18-2 | NA57-100UG | Sigma-Aldrich | 1/1000 |
| Anti-CtIP | D-4 | Sc-271339 | SantaCruz | 1/1000 |
| Anti-TRIP13 |  | A303-605A | Bethyl Laboratories | 1/1000 |
| Anti-HA |  | H6908 | Sigma-Aldrich | 1/1000 |
| Anti-NEK7 |  | A302-684A | Bethyl Laboratories | 1/1000 |
| Anti-ASC | AL177 | AG-25B-0006-C100 | AdipoGen | 1/1000 |

### TCGA data acquisition and analysis

TCGA expression data were downloaded from the UCSC Zena browser. HRD signature was analyzed according to the HR defect (HRD) score^25^. High and low HRD were defined for gene expression analyses.

### Statistical analyses

Data analyses were performed with GraphPad Prism 10.0 using the tests as indicated in the figure legends. Data are representative of at least two or three independent experiments. For each of the experiments, the type, normality and homocedasticity of the data was checked in order to choose the appropriate statistical test. Significance *p < 0.05, **p < 0.01, ***p < 0.001, ****p < 0.0001. ; ns, not significant (p ≥ 0.05).

## ACKNOWLEDGEMENTS

We are grateful to the imaging (PIC platform), flow cytometry (CYLE platform) and bioinformatics facilities (Gilles Thomas platform) of the CRCL. We thank B Manship for English editing. We are grateful to Gaëlle Legube, and Jeremy Stark, respectively, for sharing the DIvA, and U-2 OS DR-GFP cell models.

## Supplemental figures

**Supplementary Figure 1.**
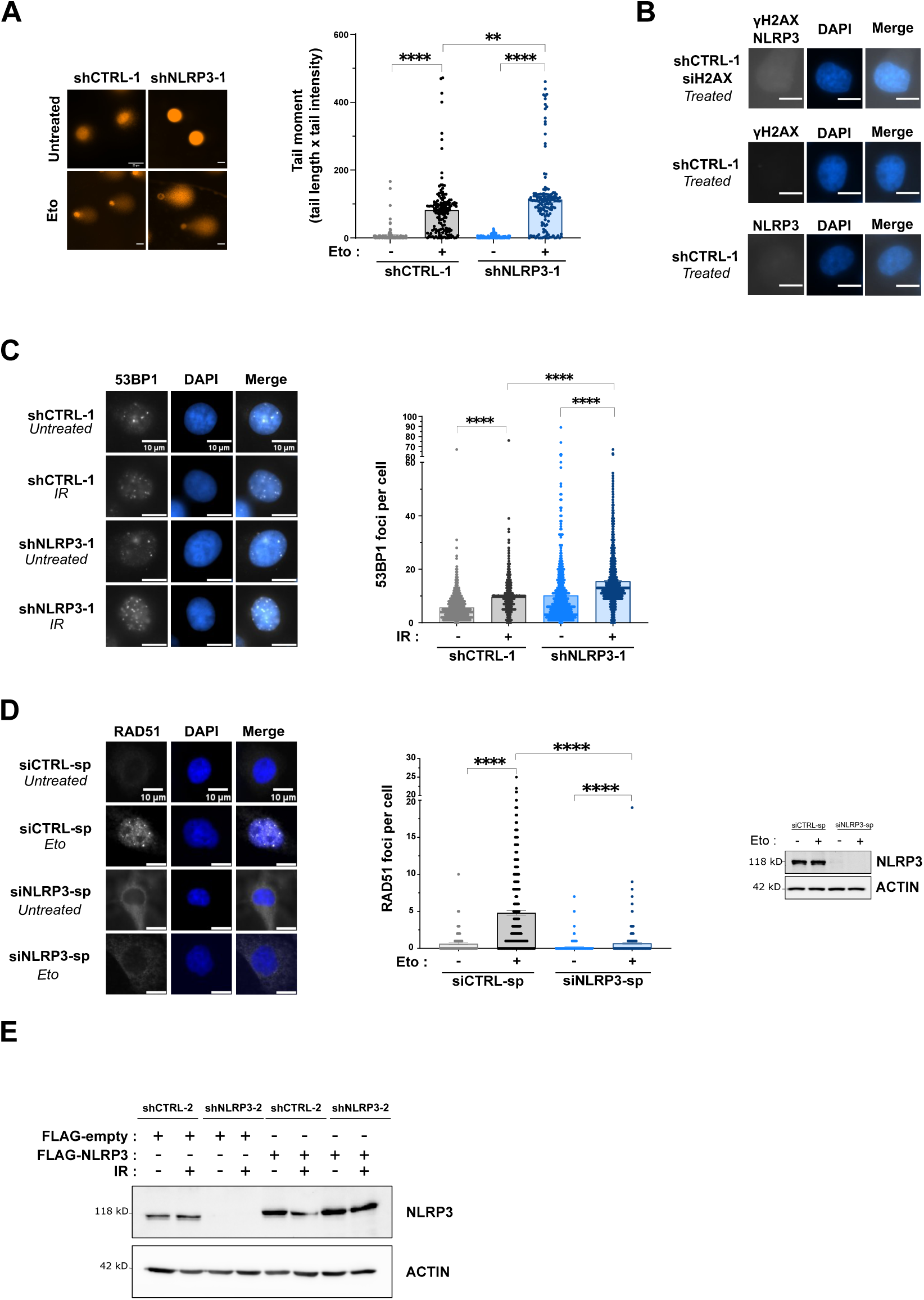
– NLRP3 facilitates homologous recombination. (A) DNA damage was analyzed by comet assay performed 1h after 100 μM etoposide treatment. Representative images of comet tails (left panel) and dot plot with comet tail moment (right panel) are shown. Representative of two independent experiments. Scale bar: 20 µm. (B) Representative microscopy images of Proximity Ligation Assay (PLA) of control conditions, in white the interaction between γH2AX and NLRP3. DAPI in blue. Scale bar: 10 µm. (C) Representative images of 53BP1 foci 1 h after 5 Gy IR treatment in MDA-MB-231 shCTRL-1 or shNLRP3-1 (left panel). DAPI was used to stain the nuclei. Scale bar, 20 μM. Quantification of 53BP1 foci is shown (right panel). Total number of foci are shown. At least 100 cells were quantified per experiment. (D) MDA-MB-231cells transfected with control (CTRL-1) or NLRP3 (NLRP3-1) siRNA were treated with 0.5 μM etoposide for 3 h, and the number of RAD51 foci was assessed by immunofluorescence (left panel). DAPI in blue. Scale bar: 10 µm. Quantification of number of cells with RAD51 foci is shown (middle panel). Western blot analysis showing NLRP3 depletion in cells 48 h post-transfection (right panel). ACTIN served as a loading control. One representative experiment of 3 biological replicates is shown. (E) Immunoblot analysis to control NLRP3 depletion and Flag-NLRP3 overexpression in MDA-MB-231 cells after irradiation. ACTIN as loading control. (F) Immunoblot analysis to control NLRP3 and CtIP depletion in U-2 OS DR-GFP cells, and gating strategy. (G) U-2 OS DR-GFP cells were co-transfected with mCherry-NLRP3 or mCherry-empty and HA-I-SceI plasmids, and the number of GFP+ cells was assessed by flow cytometry 72 h post-transfection. Two pooled experiments. (H) Sensitivity of shCTRL-1 or shNLRP3-1 MDA-MB-231 cells transfected with control (CTRL-3) or BRCA1 siRNA to indicated doses of olaparib was assessed using colony formation assay. Percentage of surviving colonies was analyzed after 14 days. (I) shRNA-mediated NLRP3 knockdown was performed in U-2 OS. U-2 OS shCTRL-1 or shNLRP3-1 were treated with indicated doses of talazoparib (Tala) and assayed for surviving colonies 14 days later. Percentage of surviving colonies was analyzed (top panel). Western blot analysis showing NLRP3 depletion in cells (bottom panel). ACTIN served as a loading control. (J) Expression of NLRP3 in primary tumors from breast cancer patients stratified by homologous recombination deficiency (HRD) low and HRD high from the TCGA cohort. HRD high, n = 525 distinct patients; HRD low, n = 525 distinct patients. The p-value correspond to a Welch Anova’s test. P value = 4.58e-15. (A,B,C,E,H,I) shRNA-mediated NLRP3 knockdown was performed in MDA-MB-231. For (D), at least 100 cells was quantified from one representative experiment. Mean and SEM for at least two (A,C,G) or three (H,I) independent replicas is shown. (A,J) The p-value correspond to an unpaired T-test with Welch’s correction. (C,D,H,I) The p-values correspond to a Mann Whitney’s test. ns: nonsignificant ** p < 0.01, **** p < 0.0001.

**Supplementary Figure 2.**
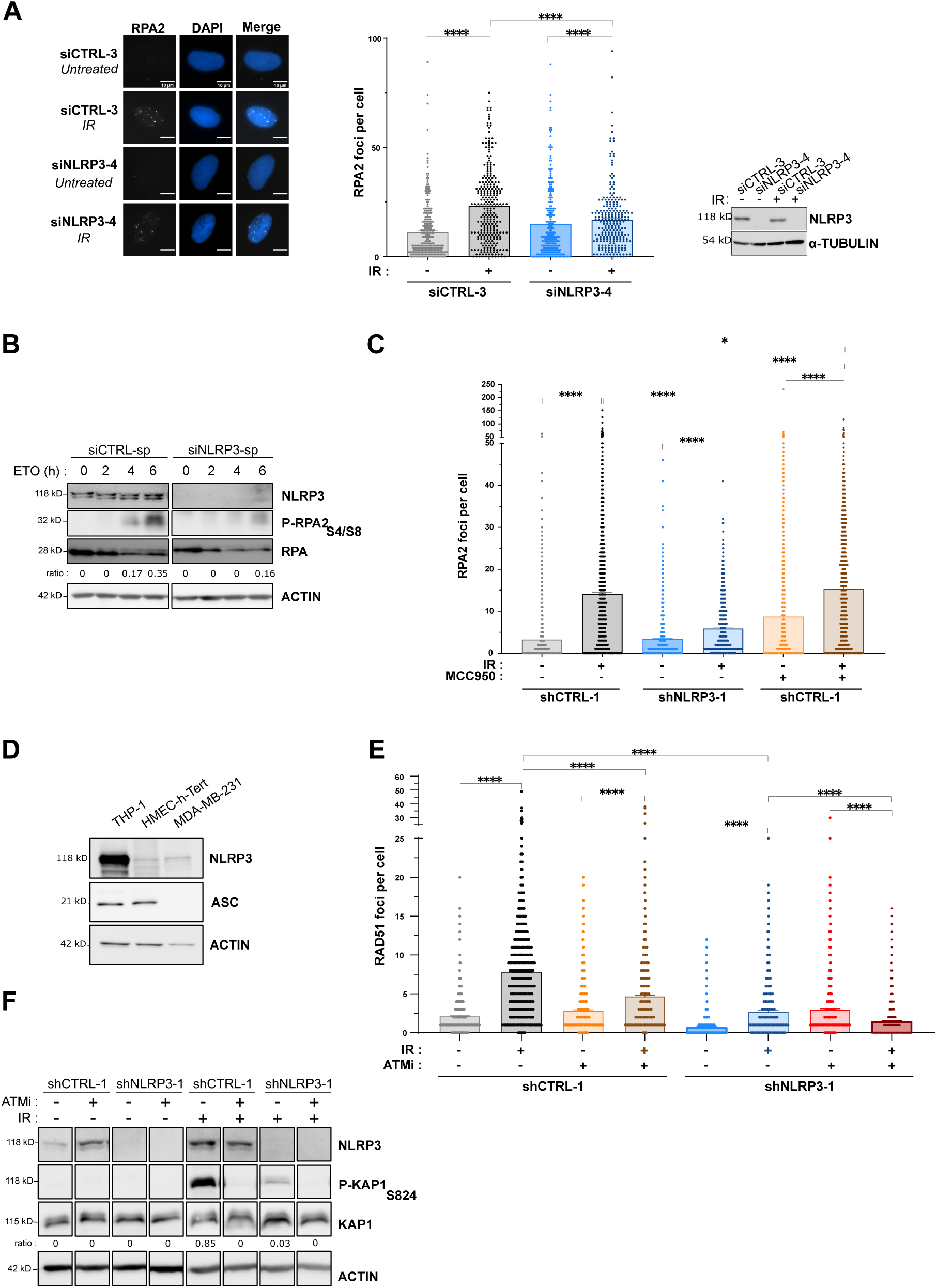
– NLRP3 is required for DNA end resection at DSB. **(A)** RPA2-foci in U-2 OS transfected with CTRL-3 or NLRP3-4 siRNA and treated with IR (5 Gy) for 6 h. Cells were selected with blasticidin. Representative images of RPA2 foci (left panel) and dot plot with RPA2 foci per cells are shown (middle panel). Scale bar: 10 µm. Control immunoblot for knock-down efficacy in U-2 OS (right panel). TUBULIN served as a loading control. Dots represent n=2 biological replicates, 200-300 cells analyzed, mean ± SEM, Mann Whitney test. * p < 0.05, **** p < 0.0001. **(B)** MDA-MB-231 transfected with siCTRL-sp or siNLRP3-sp were treated with 100 μM etoposide. Phospho-RPA2 (S4/S8) (pRPA2) was monitored over time by immunoblot. The ratio in relative units of pRPA2/RPA2 are shown. ACTIN served as a loading control. Representative of 2 independent experiments. **(C)** Quantification of RPA2 foci in shCTRL-1 or shNLRP3-1 MDA-MB-231 cells after IR in the presence of DMSO (-) or NLRP3 inhibitor (MCC950) (+) (5 Gy, 6 h). Dot plot with total number of foci per cell are shown. **(D)** Control immunoblot in THP-1, HMEC-h-Tert and MDA-MB-231 cells for NLRP3 and ASC expressions. ACTIN as loading control. Representative of 2 independent experiments. **(E)** Quantification of RAD51 foci in shCTRL-1 or shNLRP3-1 MDA-MB-231 cells after IR in the presence of DMSO (-) or ATM inhibitor (KU55933) (+) (5 Gy, 6 h). Dot plot with total number of foci per cell are shown. **(F)** Immunoblot analysis to control NLRP3 depletion and treatment with ATMi in MDA-MB-231 cells after irradiation. The efficacy of the treatment was demonstrated by the reduction in phosphorylation of an ATM effector, KAP1. The ratio in relative units of P-KAP1/KAP1 are shown. ACTIN as loading control. (C, E, F) shRNA-mediated NLRP3 knockdown was performed in MDA-MB-231. (A-C-E) Total number of foci are shown. At least 100 cells were quantified per experiment. Mean and SEM of 2 (A, C, E) independent experiments. (A, C, E) The p-value correspond to a Mann Whitney’s test. ** p < 0.01, **** p < 0.0001.

**Supplementary Figure 3.**
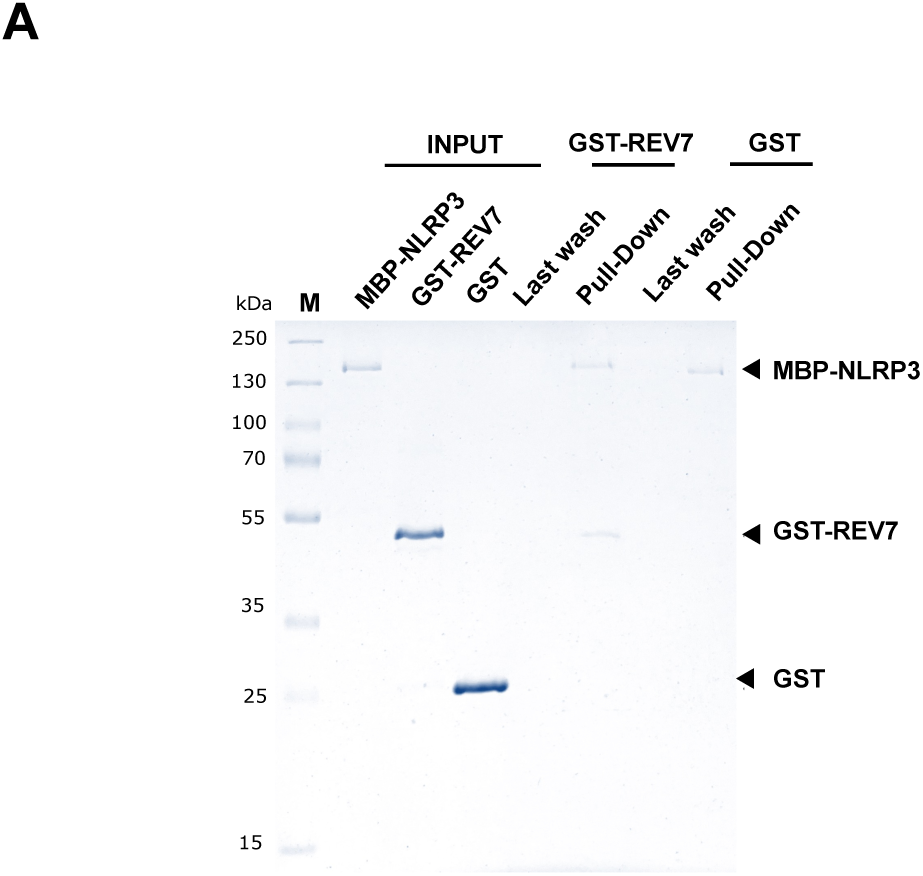
– NLRP3 interaction with REV7 is direct. (A) Pull down experiment of separately expressed and purified NLRP3 and REV7 proteins. Recombinant MBP-NLRP3, GST-REV7 and GST proteins are shown in input lanes 1-3. Pull down of GST-REV7 with MBP-NLRP3 as bait indicates a direct interaction (lane 5), whereas GST alone is not binding (lane 7). The supernatant after three washing steps is shown as a control (lanes 4 and 6). One representative out of 3 experiments.

**Supplementary Figure 4.**
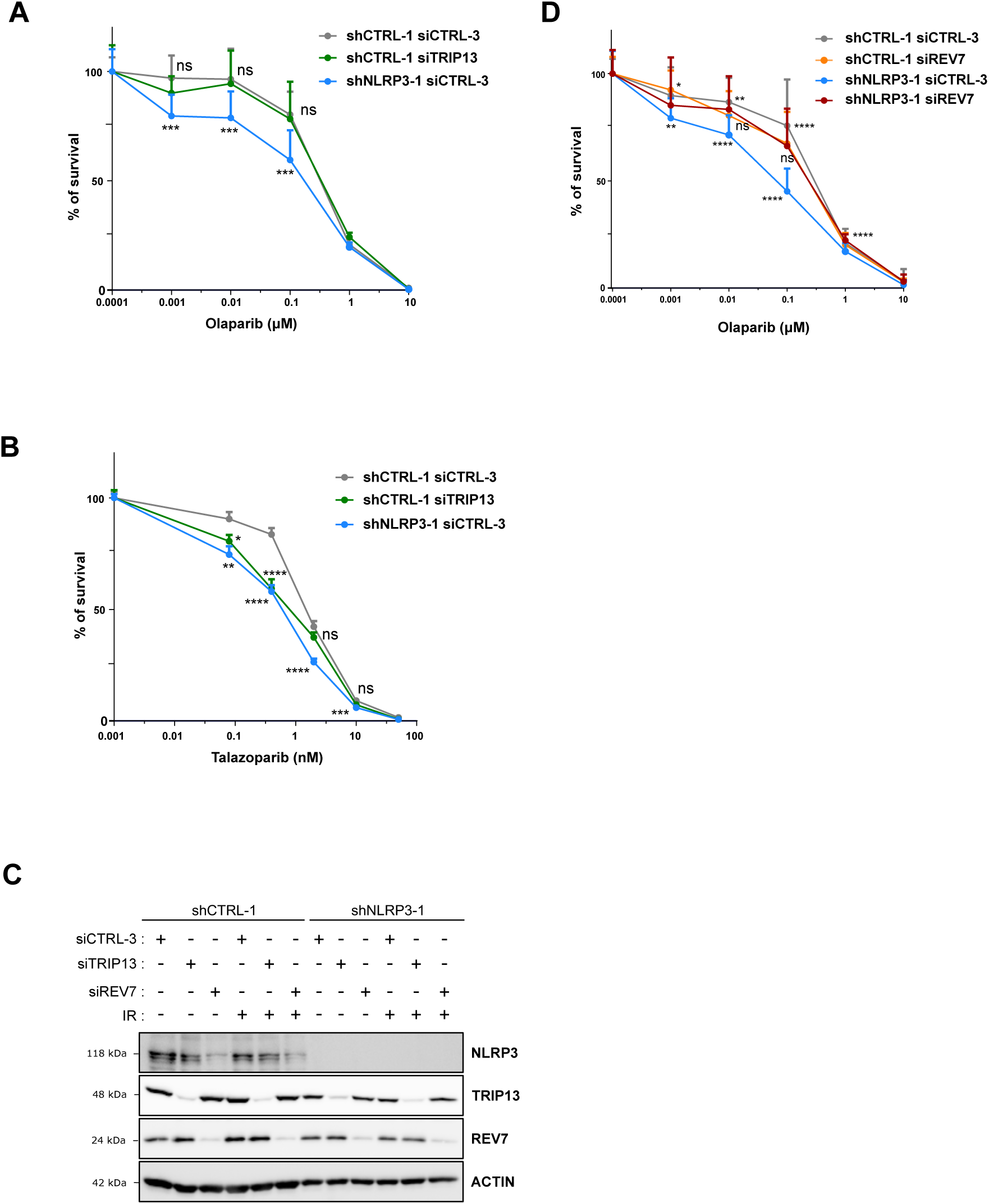
– Loss of REV7 abolishes PARPi sensitivity of NLRP3-depleted cells. (A-D) shRNA-mediated NLRP3 knockdown was performed in MDA-MB-231. MDA-MB-231 shCTRL-1 or shNLRP3-1 were treated with CTRL-3, TRIP13 or REV7 siRNA, then with indicated doses of olaparib (A,D) or talazoparib (B) and assayed for using CFA. (A-B) The same data were used for the shNLRP3-1 siCTRL-3 and shCTRL siCTRL-3 conditions as in Figure 1F, 4B and Sup 1G but the shCTRL-1 siTRIP13 data were implemented. (C) Immunoblot analysis to control NLRP3, TRIP13 and REV7 depletions in MDA-MB-231 cells after irradiation. ACTIN served as a loading control. (A,B,D) Mean and SEM for at least three independent replicas is shown. The p-value correspond to a Mann Whitney’s test. ns: nonsignificant, * p < 0.05, ** p < 0.01, *** p < 0.001, **** p < 0.0001.

**Supplementary Figure 5.**
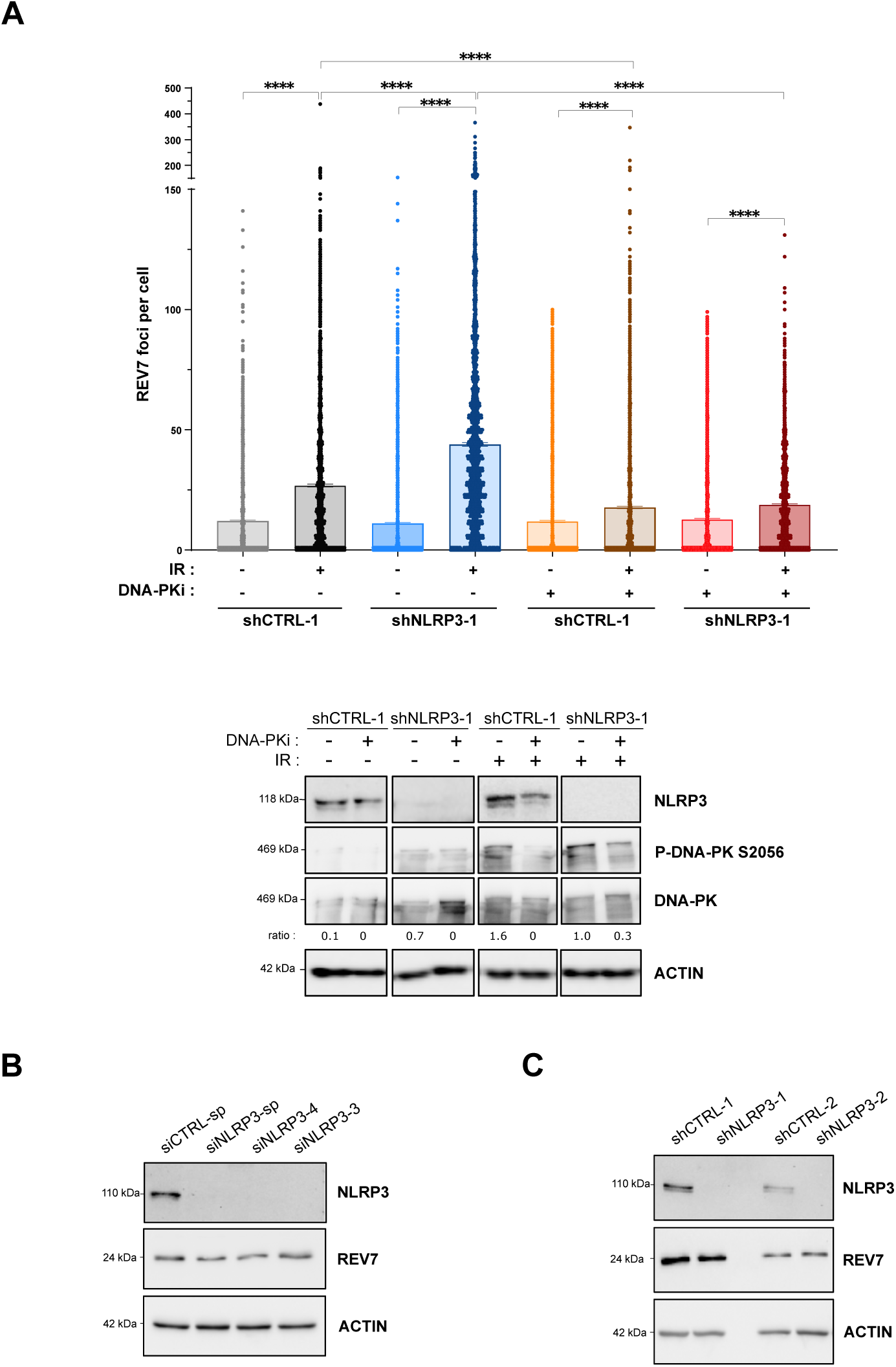
– NLRP3 inhibits REV7 recruitment to DSB. **(A)** shCTRL-1 or shNLRP3-1 MDA-MB-231 cells were pre-treated with DNAPK inhibitor NU7441 (DNAPKI) (+) or DMSO (-) then treated with IR (top panel). REV7 foci were analyzed 6 h post-IR (5 Gy). At least 100 cells were quantified per experiment. Mean and SEM for three independent replicas is shown. The p-values correspond to a Mann Whitney’s test. ns: nonsignificant, **** p < 0.0001. Immunoblot analysis to control NLRP3 depletion and treatment with DNA-PKi in MDA-MB-231 cells after irradiation (bottom panel). The efficacy of the treatment was demonstrated by the reduction in phosphorylation of DNA-PK. The ratio in relative units of pDNA-PK/DNA-PK are shown. ACTIN as loading control. **(B)** Immunoblot demonstrating that loss of NLRP3 expression induced by different siRNA does not alter the protein level of REV7 in MDA-MB-231 cell line. ACTIN was used as a loading control. Representative of at least 3 independent experiments. **(C)** Immunoblot showing that REV7 protein level is not affected in shNLRP3-1 or shNLRP3-2 MDA-MB-231 cell lines compared to control cells. Representative of at least 3 independent experiments. (A,C) Knockdown of NLRP3 using shNLRP3-1 or shNLRP3-2 was performed in MDA-MB-231.

